# Toward *in vivo*-relevant hERG safety assessment and mitigation strategies based on relationships between non-equilibrium blocker binding, three-dimensional channel-blocker interactions, dynamic occupancy, dynamic exposure, and cellular arrhythmia

**DOI:** 10.1101/2020.06.08.139899

**Authors:** Hongbin Wan, Gianluca Selvaggio, Robert A. Pearlstein

## Abstract

The human ether-a-go-go-related voltage-gated cardiac ion channel (commonly known as hERG) conducts the rapid outward repolarizing potassium current in cardiomyocytes (I_Kr_). Inadvertent blockade of this channel by drug-like molecules represents a key challenge in pharmaceutical R&D due to frequent overlap between the structure-activity relationships of hERG and many primary targets. Building on our previous work, together with recent cryo-EM structures of hERG, we set about to better understand the energetic and structural basis of promiscuous blocker-hERG binding in the context of Biodynamics theory. We propose a two-step blocker binding process consisting of:

1. Diffusion of a single fully solvated blocker copy into a large cavity lined by the intracellular cyclic nucleotide binding homology domain (the initial capture step). Occupation of this cavity is a necessary but insufficient condition for ion current disruption.
2. Translocation of the captured blocker along the channel axis (the I_Kr_ disruption step), such that:

a. The head group, consisting of a quasi-linear moiety, projects into the open pore, accompanied by partial de-solvation of the binding interface.
b. One tail moiety packs along a kink between the S6 helix and proximal C-linker helix adjacent to the intra-cellular entrance of the pore, likewise accompanied by mutual de-solvation of the binding interface (noting that the association barrier is comprised largely of the total head + tail group de-solvation cost).
c. Blockers containing a highly planar moiety that projects into a putative constriction zone within the <u>closed</u> channel become trapped upon closing, as do blockers terminating prior to this region.
d. A single captured blocker molecule may associate and dissociate from the pore many times before exiting the CNBHD cavity.

Lastly, we highlight possible flaws in the current hERG safety index (SI) and propose an alternate *in vivo-relevant* strategy factoring in:

1. Benefit/risk.
2. The predicted arrhythmogenic fractional hERG occupancy (based on action potential simulations of the <u>undiseased</u> human ventricular cardiomyocyte).
3. Alteration of the safety threshold due to underlying disease.
4. Risk of exposure escalation toward the predicted arrhythmic limit due to patient-to-patient pharmacokinetic variability, drug-drug interactions, overdose, and use for off-label indications in which the hERG safety parameters may differ from their on-label counterparts.

## Introduction

As is widely appreciated throughout the pharmaceutical industry, the risk of torsade de pointes arrhythmia (TdP) is promoted by drug-induced loss of function of the human ether-a-go-go gene product (hERG) [1–4] in proportion to the concomitant fractional decrease in the outward repolarizing hERG current (denoted I_Kr_) due to dynamic blocker occupancy. TdP arises at a threshold level of I_Kr_ reduction, which may auto-extinguish spontaneously or progress to ventricular fibrillation and death [5–8]. The causal relationship between acquired loss of hERG function and TdP was deduced in the late 1990s, resulting in the withdrawal or black box labeling of several implicated drugs [9,10], together with the implementation of routine hERG safety assessment and monitoring practices during the preclinical and clinical stages of pharmaceutical R&D. A surprisingly high prevalence of hERG blockade throughout drug-like chemical space was revealed in the process, the molecular causes of which have been investigated with limited success over the years using a number of experimental and *in silico* approaches. Inadvertent hERG blockade arises frequently among screening hits and early leads, which is often reduced but rarely eliminated via trial-and-error chemical optimization. As a result, hERG activity remains a highly problematic liability during both the preclinical and clinical stages of drug R&D. The possibility that hERG blockade is only one part of a multi-dimensional ion channel safety problem has been raised by the Comprehensive *in vitro* Pro-arrhythmia Assay (CIPA) initiative [11]. However, because hERG blockade is far more prevalent than that of other cation channels, multi-channel blockade likely accounts for only a subset of TdP cases (TdP is evoked when the inward-outward current balance is tipped toward the inward direction beyond a threshold level, irrespective of the cause).

Fractional I_Kr_ reduction manifests as a graded prolongation of the ventricular action potential duration (APD), mirrored by the QT interval in the electrocardiogram (ECG). Late-stage preclinical hERG safety assessment consists of rising dose ECG studies in dogs or primates aimed at determining the no observed adverse effect QT prolongation (LQT) exposure level, which is necessarily far above the projected human therapeutic free plasma C_max_ (hereinafter referred to as TFPC_max_).

Preclinical hERG safety assessment and mitigation are chicken-egg problems, in which either the maximum safe therapeutic free plasma C_max_ is limited by the maximum achievable hERG IC_50_, or the minimum safe hERG IC_50_ is limited by the minimum efficacious free plasma C_max_. The idea is to maintain a safe “distance” between hERG occupancy at the therapeutic and arrhythmic exposure levels, allowing for unintended exposure-driven occupancy escalation in the patient population. However, the status quo hERG safety index (SI) [12] is based on an entirely empirical model (expressed as the maximum safe free plasma C_max_ ≤ 1/30 · hERG IC_50_) that we show in this work possibly suffers from multiple flawed assumptions.

*In vitro* hERG potency is necessarily mitigated to the lowest possible level (ideally, the limit of detection), constrained by primary target potency, solubility, permeability, and many other requisite drug-like properties. Mitigation is typically approached via trial-and-error chemical analoging guided by *in vitro* testing and *in silico* prediction, the objective of which is to achieve a therapeutic index (TI) *in vivo* (the ratio of the TFPC_max_ to the arrhythmic C_max_ or a designated LQT threshold). However, the hERG TI in humans cannot be predicted reliably in the absence of human pharmacokinetic (PK) data, including the TFPC_max_ and extent of exposure escalation resulting from patient-to-patient PK variability, drug-drug interactions (DDI), and/or overdose. Instead, preclinical safety assessment is performed using safety indices (SI) based on general off-target potency/exposure relationships. Redfern et al. developed the following hERG SI based on potency, clinical PK data (including the highest reported C_max_), and reported TdP cases for 100 marketed drugs (including anti-arrhythmic hERG blockers), ranging from no reported cases to TdP-linked withdrawals [12]:

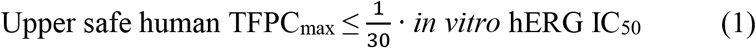

The Redfern SI implicitly accounts for exposure escalation via a wide 30-fold margin between the *in vitro* hERG IC_50_ and TFPC_max_, which as we demonstrate below, translates to effectively zero tolerated hERG <u>occupancy</u> at the maximum anticipated therapeutic exposure in humans. However, the safety margin for compounds exhibiting residual hERG activity at the projected TFPC_max_ cannot be assessed systematically or tailored to benefit/risk via the all-or-none Redfern criterion. Considerable time and effort may be invested in hERG mitigation to the limit of detection, which is subject to the following caveats:

1. Constraints on chemical mitigation imposed by the typically high overlap among the structural and physico-chemical properties promoting hERG and primary target potency, solubility, permeability, and PK behaviors.
2. Insufficient accuracy of *in vitro* assays (typically, radio-ligand displacement and automated or manual patch clamp) needed to resolve true hERG structure-activity relationships (SAR). *In vitro* hERG IC_50_ values were shown to vary as a function of cell culture conditions, patch clamp protocol, data fitting approach [13–15], and temperature [14]. IC_50_ variation of 16- and 23-fold has been reported for terfenadine and loratadine, respectively
3. The limited relevance of *in vitro* hERG binding/blockade measurements to <u>none-quilibrium</u> conditions *in vivo.*
4. The lack of *in vivo* pro-arrhythmia assessment during the lead optimization stage, which is typically reserved for late-stage clinical candidates (relegating the SI to a prediction of *in vivo* behavior).

In our previous works:

1. We studied dynamic hERG blockade by compounds that are trapped within closed channels (“trappable” blockers) and those that are expelled during closing (“non-trappable” blockers) using a version of the O’Hara-Rudy model of the <u>undiseased</u> human ventricular cardiomyocyte [16] into which we introduced a Markov hERG blocker binding schema [17]. We showed that blockade by non-trappable blockers builds and decays in tandem with channel opening and closing, respectively, whereas trappable blocker occupancy accumulates with increasing exposure up to the free C_max_.
2. We derived a general analytical treatment of non-equilibrium binding that accounts for binding site buildup and decay cycles driven by translocation, conformational changes, or synthesis and degradation of the binding partners [18]. We showed that binding under dynamic conditions is characterized by time-dependent occupancy (rather than static occupancy/potency), which is governed by k_on_ relative to the rate of binding site buildup, concentration/exposure, and k_off_ relative to the rate of binding site decay. Non-trappable hERG blocker binding clearly falls at the extreme end of the non-equilibrium spectrum, given that the open (blocker-accessible) state of hERG normally builds <u>and</u> decays over a 350-400 ms time window [17].
3. We hypothesized that hERG binding is energetically driven by de-solvation and resolvation costs (the putative origin of all non-covalent binding free energy barriers [17,19–23]). Toward that end, we studied the solvation properties of the hERG pore using WaterMap and a homology model of the protein [24] (prior to the publication of the cryoEM structure of open state hERG). The results of our calculations suggest that the pore lumen is solvated almost exclusively by bulk-like and hydrogen bond (H-bond) depleted water, consistent with low blocker association cost (i.e. no or low de-solvation cost of the pore) and high blocker dissociation cost (i.e. high re-solvation cost of the pore). Association and dissociation costs are therefore plausibly relegated largely to <u>blocker desolvation</u> and <u>pore re-solvation</u> costs, respectively [17]. Additionally, the association rate is plausibly enhanced by electrostatic interactions between basic blockers and the negative field within the pore.
4. We showed that k_off_ of <u>non-trappable</u> blockers from static channels that is slower than the closing rate in dynamic channels is “hijacked” by the closing rate. Since hERG patch clamp assays are typically run at sub-physiological gating frequencies (and gating is neglected altogether in radio-ligand binding assays), the hijacking effect is likely underestimated to varying degrees. The rate of channel closing dominates over the dissociating pore and blocker re-solvation costs.

Here, we use Biodynamics principles [18] to examine the structural basis of hERG blockade (including trappability) at the atomistic level, and revisit hERG safety assessment in an *in vivo*-relevant context.

## Materials and methods

All calculations and visualizations were performed using Maestro 2019-1 (Schrodinger, LLC, Portland, OR) on a representative set of canonical hERG blockers taken from reference [12] (Table 1), as well as trappable and non-trappable propafenone analogs taken from reference [25] (Table 2). The structures were built using LigPrep and energy minimized using Macromodel (MMFF force-field, and default parameters). The hERG structure (PDB code 5VA2) [26] was prepared (hydrogen addition, Asn, Gln, His orientations) using PPrep. Blocker docking sites and the solvation properties thereof were characterized using SiteMap. Overlay models were generated via manual superposition on a template consisting of the GDN detergent crystallized with Na_v_1.4 (PDB code 6AGF) [27], which we propose as the general intra-cellular binding mode for voltagegated ion channel blockers. We emphasize the theory/knowledge-, rather than computation-driven, underpinnings of this work, the core tenets of which include: 1) the dependence of dynamic occupancy on binding partner buildup and decay under non-equilibrium conditions *in vivo*, wherein the applicability of equilibrium binding metrics (e.g. ΔG, K_d_, K_i_, IC_50_, EC_50_) is limited primarily to the *in vitro* setting [17,18]; and 2) the predominant contribution of de-solvation and re-solvation costs to non-covalent kinetic barriers under aqueous conditions (a core tenet of Biodynamics theory [18,23]).

**Table 1.**
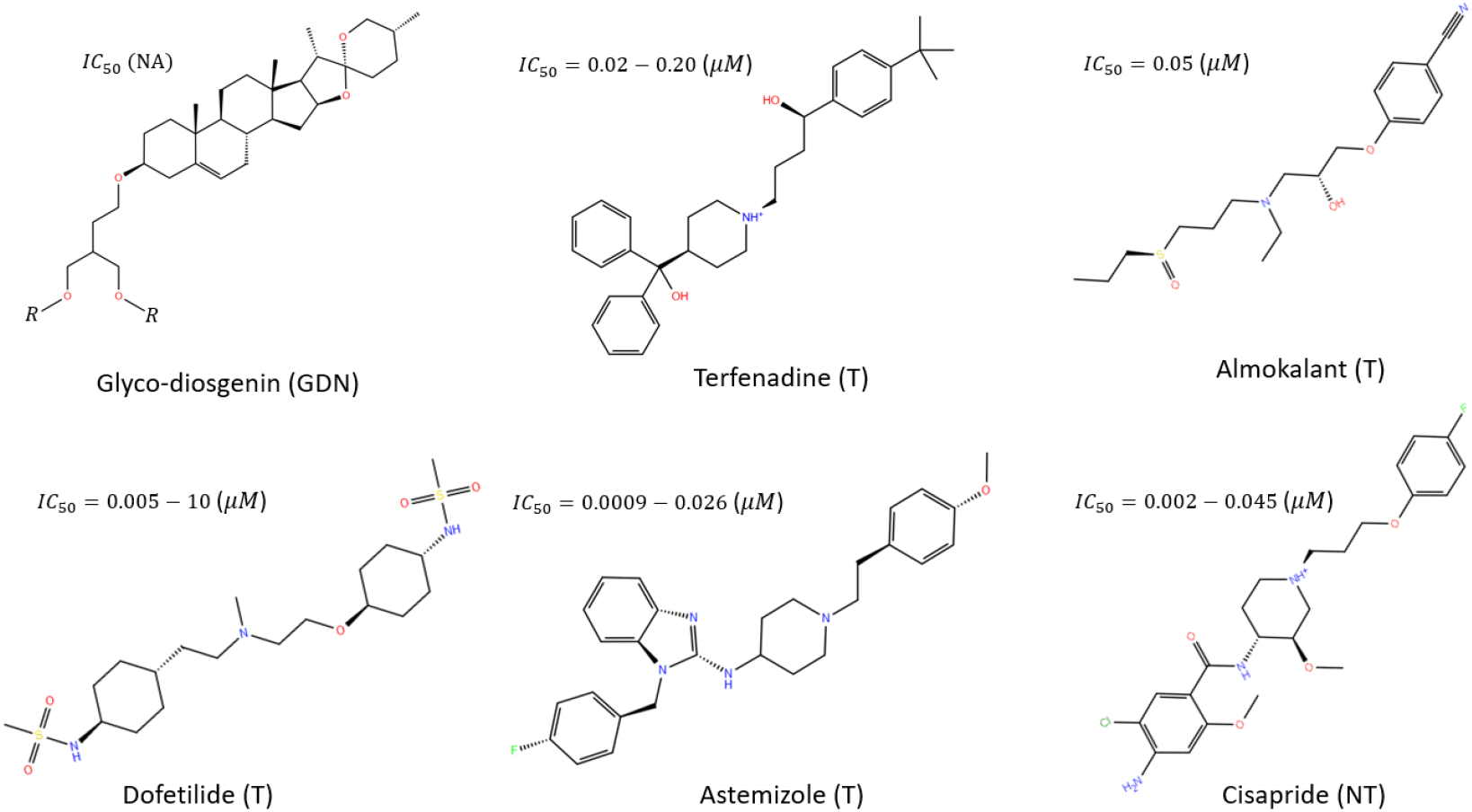
Representative hERG blockers studied in this work [12].

**Table 2.**
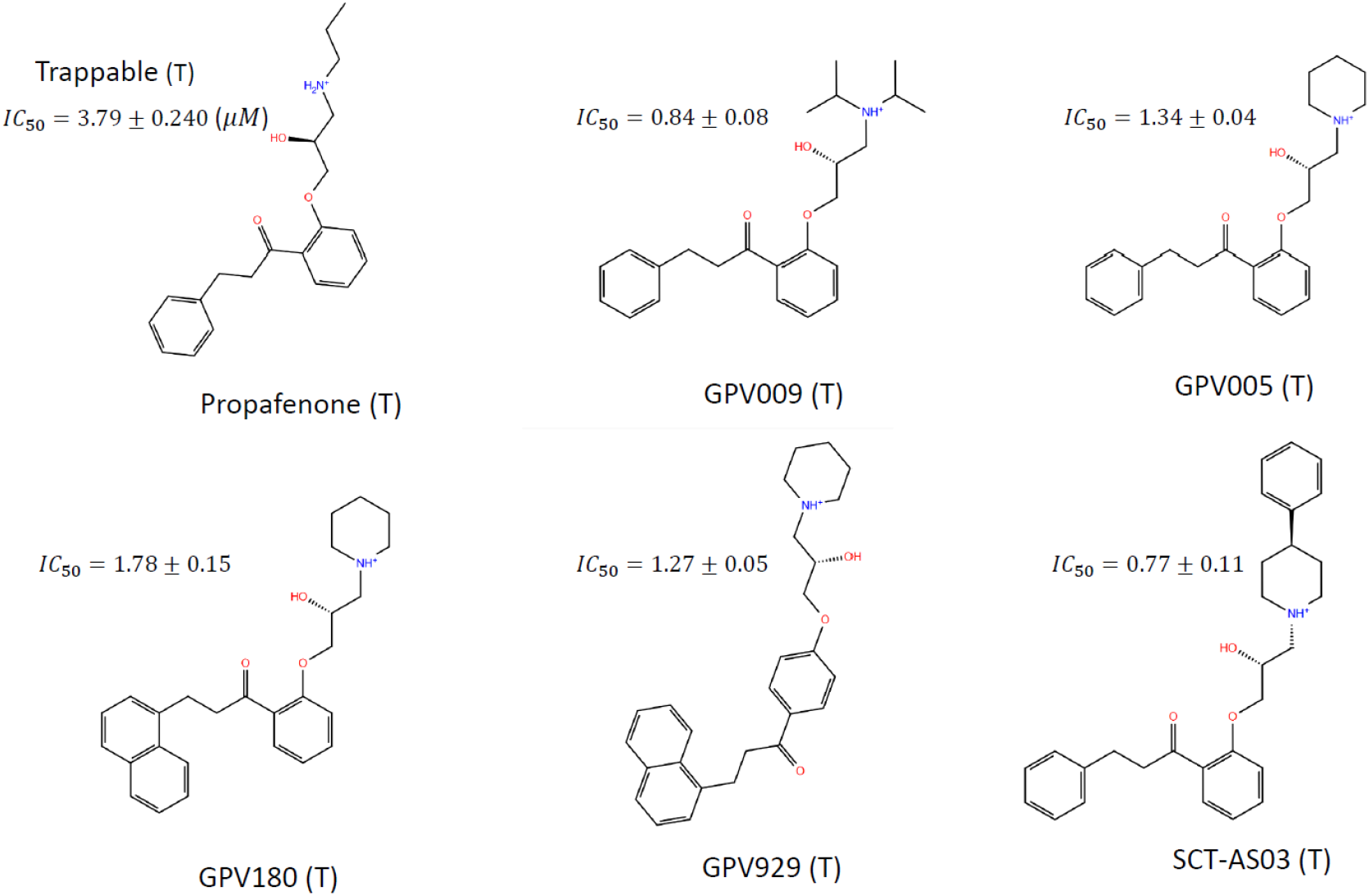
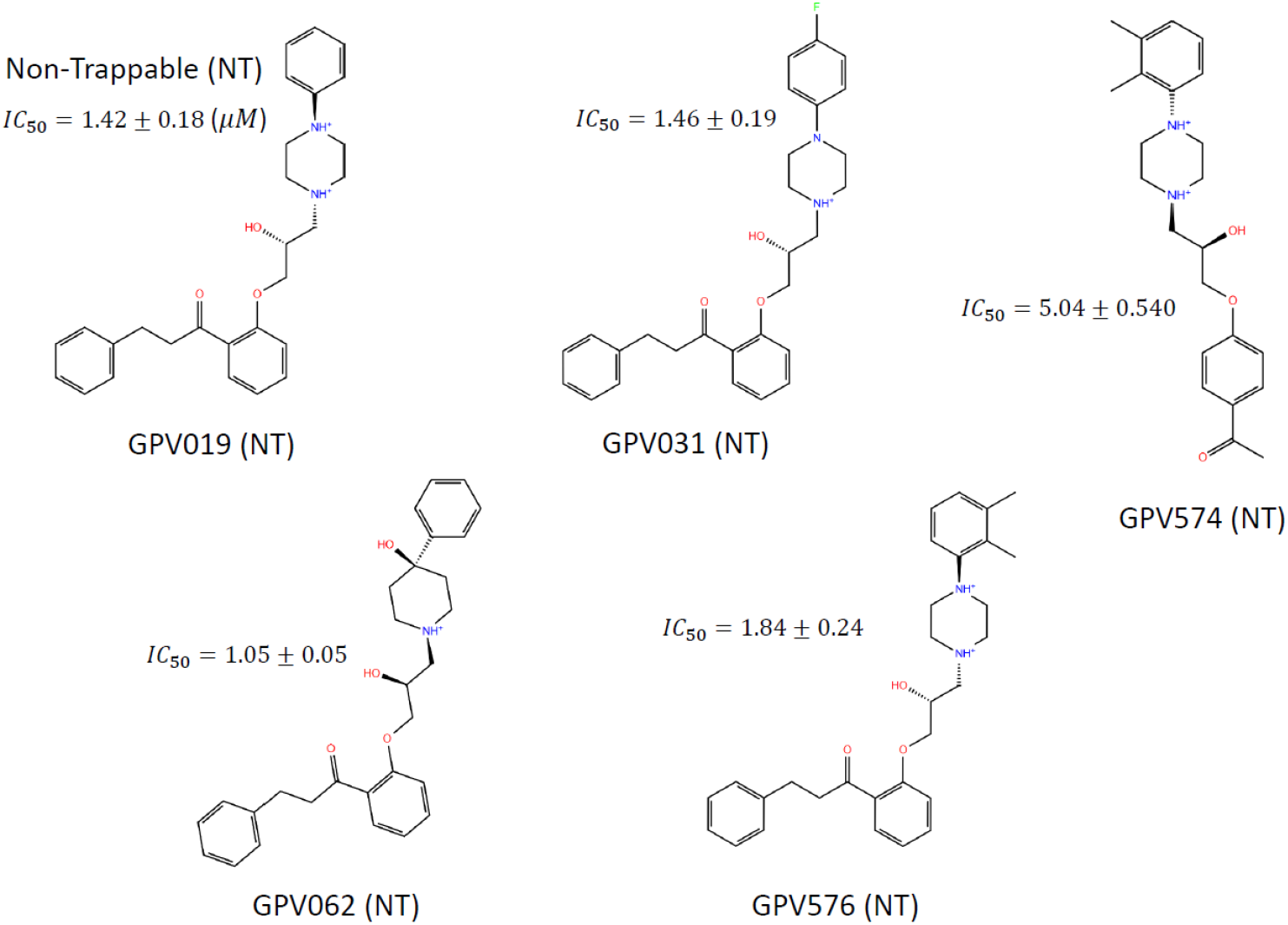
Published trappable and non-trappable hERG blockers studied in this work [25].

## Results

We characterized the energetic, structural, and chemical drivers of blocker binding and trappability using modeled structures of known trappable and non-trappable compounds, together with a set of recently published cryo-EM structures of the full length open hERG channel (PDB codes 5VA1, 5VA2, 5VA3) [26]) [26]. Our overall findings suggest that blocker binding is governed by the following contributions:

1. Steric shape and size complementarity between blockers and the pore binding region.
2. Blocker de-solvation rate (proportional to the de-solvation free energy cost) vis-à-vis the channel-opening rate, together with pore and blocker re-solvation rates (proportional to the re-solvation free energy cost) vis-à-vis the channel-closing rate.
3. Blocker basicity/pKa vis-à-vis the negative field within the pore, which speeds the association rate.

Next, we outline a more *in vivo*-relevant hERG <u>mitigation strategy</u> based on these findings. Lastly, we revisit the status quo hERG <u>safety assessment protocol</u>, and propose a more *in vivo*-relevant strategy centered on the putative relationship between dynamic occupancy, PK, and cellular arrhythmogenesis.

### Blockade-relevant aspects of hERG structure and function

The hERG channel is a tetrameric protein comprised principally of Per-Arnt-Sim (PAS), transmembrane voltage sensing, transmembrane pore, C-linker, and intra-cellular cyclic nucleotide binding homology (CNBH) domains (one per monomer) (Figure 1). We speculate that the CNBH domain serves to bind the independent tetrameric subunits together via the expulsion of intersubunit H-bond depleted solvation (noting that the absence of this domain in Na_v_1.5 and Ca_v_1.2 is consistent with the uni-chain composition of those channels). A large intra-cellular tunnel-like cavity within the combined CNBH/C-linker domains (which we refer to as the “outer vestibule”) is observed in the cryo-EM structure, through which blockers must necessarily transit to reach the pore region.

**Figure 1.**
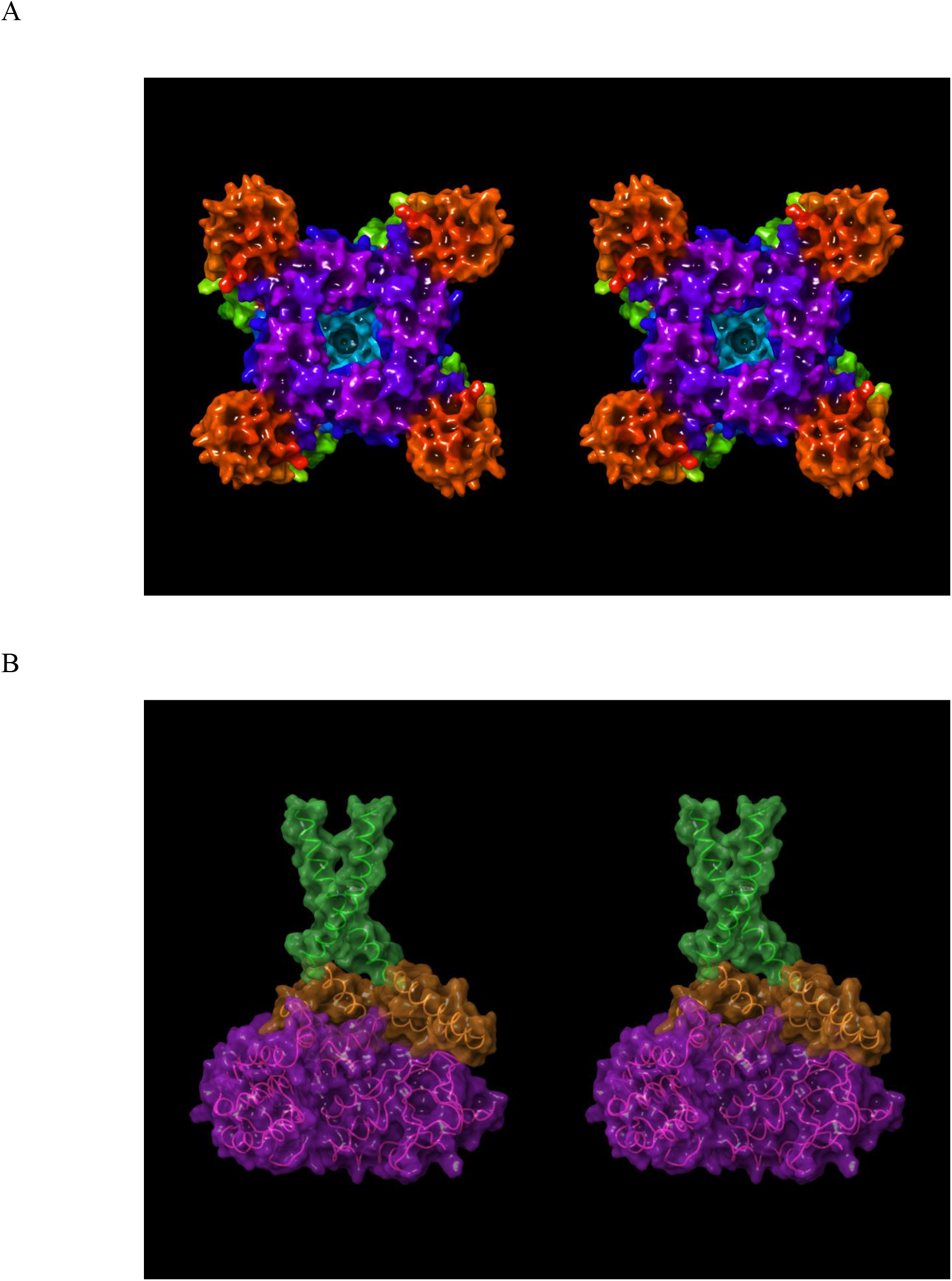
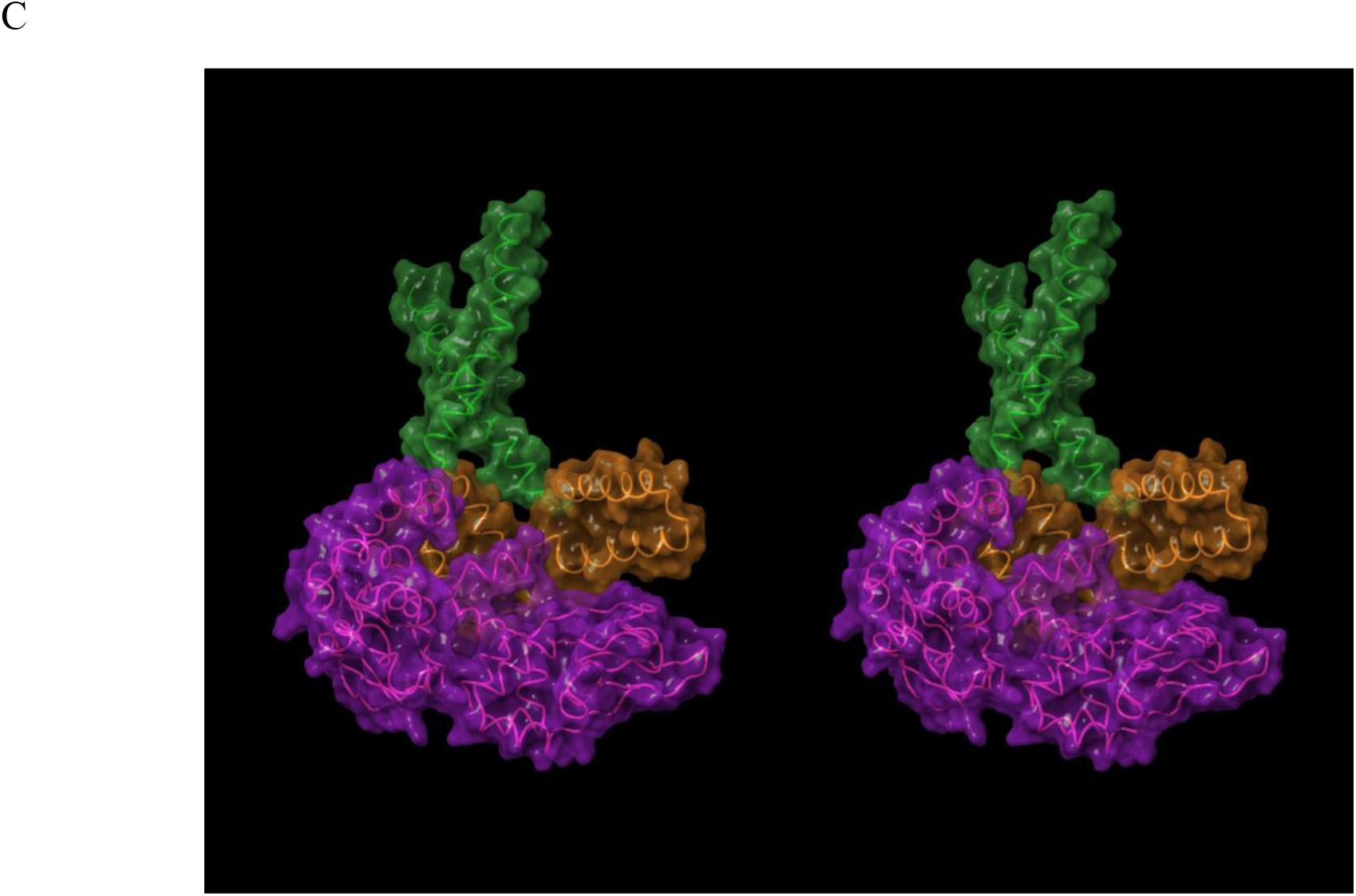
(A) Stereo image of the open state hERG cryo-EM structure (PDB code 5VA1 [26]) viewed along the pore axis from the intra-to extra-cellular direction. The pore (highlighted in cyan) and CNBH domain (highlighted in purple) cavities reside at the distal and proximal ends of the structure, respectively. The helical C-linker domains reside at the four corners of the channel (highlighted in orange), with the voltage-sensing domains (highlighted in light green) partitioned above in the membrane. The C-linker domains reside between the pore and CNBH domains, forming a continuous cavity with the latter (which we refer to as the “outer vestibule”). The PAS domain and linker were omitted from the crystallized protein construct. (B-C) Longitudinal cutaway views of the intra-cellular region of the ion conduction pathway, consisting of the Clinker lined cavity (highlighted in orange), which is sandwiched between the intra-cellular pore entrance (highlighted in green) and CNBH domain cavity (highlighted in magenta).

### The predicted canonical binding mode of voltage-gated ion channel blockers

It is widely assumed that bound hERG blockers are fully buried within the pore domain based on mutagenesis data and *in silico* docking [30,31], which we had likewise assumed in our previous work [24]. However, this hypothesis is inconsistent with the large size/volume of many blockers, wherein binding would likely depend on an unreasonably high level of induced fit. A glyco-diosgenin (GDN) detergent molecule (Figure 2A) bound to the closed state of the human voltagegated Na_v_1.4 channel observed in a recent cryo-EM structure (PDB code 6AGF) [27] offers a possible clue as to the general binding mode of cation channel blockers. GDN straddles the pore and pore entrance with its rigid polycyclic moiety buried within, and its two unresolved disaccharide moieties projecting out to the cytoplasm (reminiscent of a drain-plug) (Figure 2B-C). We proceeded to test whether hERG blockers could potentially bind in a similar “drain-plug-like” fashion using a ligand-based overlay model (noting that the cytoplasmic hERG blocker moiety would necessarily reside within the outer vestibule).

**Figure 2.**
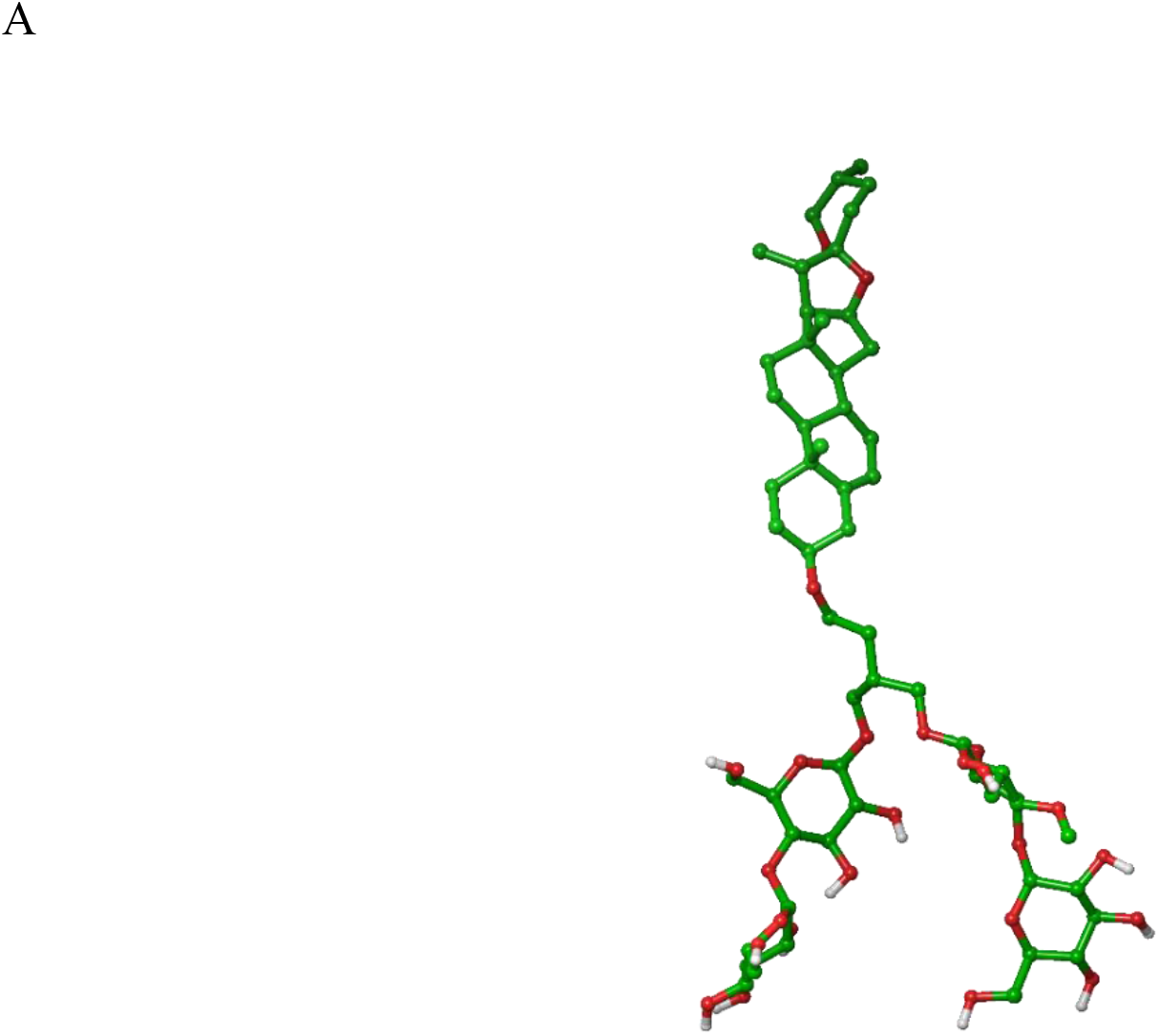
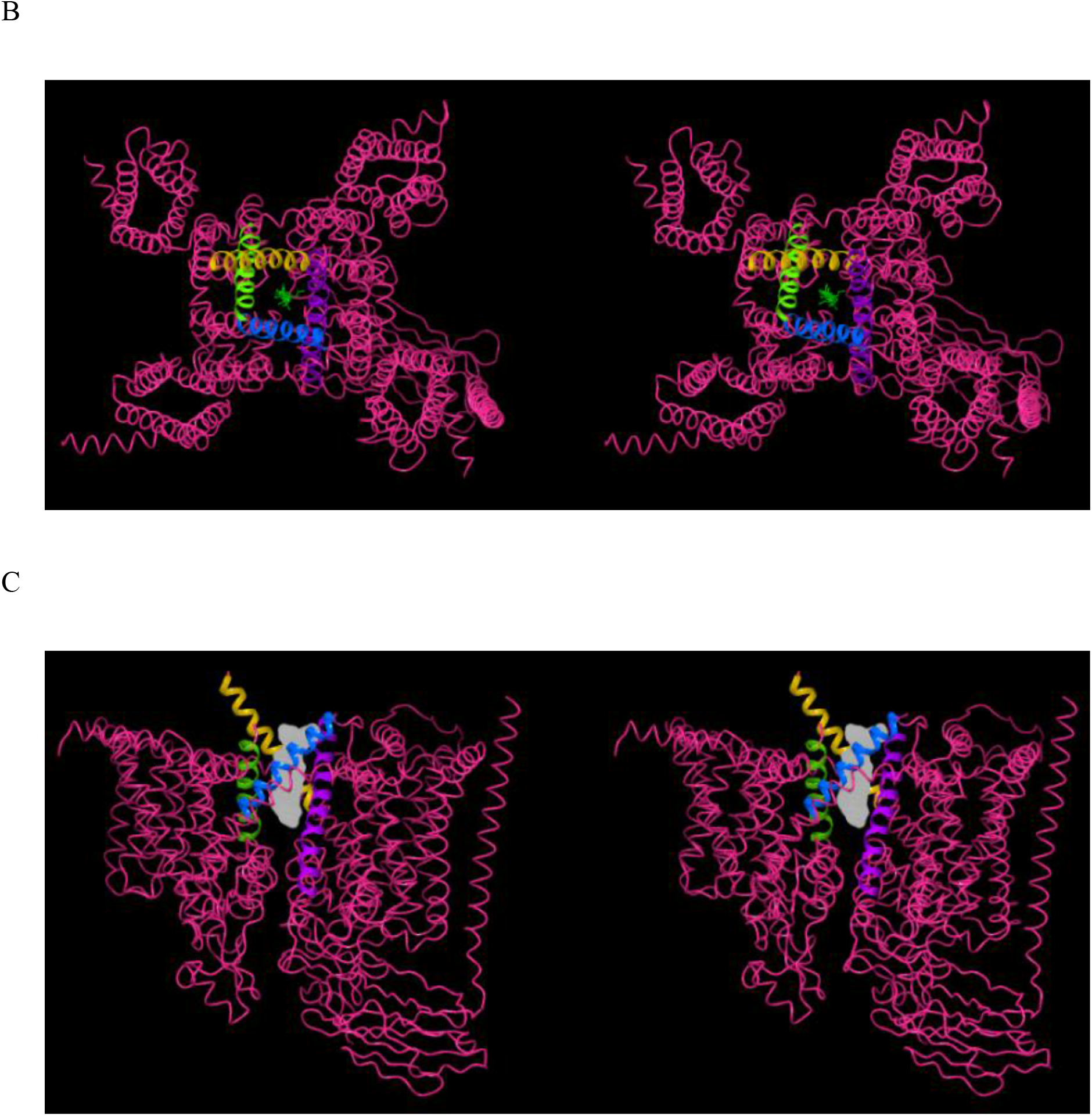
(A) The structure of GDN (PDB code 6AGF [27]), with the unresolved disaccharide moieties residing at the bottom of the structure qualitatively modeled in. (B) The Na_v_1.4 channel viewed parallel to the pore axis from the intra-to extra-cellular direction, showing the GDN molecule (green) bridging between the pore (which is partially closed) and cytoplasm (corresponding to the C-linker-enclosed cavity in hERG). (C) Same as A, except viewed perpendicular to the pore axis, with the cytoplasmic end of the channel at the top of the figure. The molecular surface of GDN is shown in gray.

We overlaid the hERG and Na_v_1.4 structures, and fit our reference set of hERG blockers (see Materials and methods) to the common pore-bound moiety of GDN (Figure 3). According to our model, the typically Y-or L-shaped hERG blockers project a single quasi-linear/rod-shaped moiety into the pore. We assume that basic nitrogen-containing moieties, when present, reside within this region. Total burial within the pore is unlikely (other than relatively small quaternary alkylamines), given the diverse range of size, chemical composition, and structural variability of the known blocker chemical space. Our analysis suggests the existence of three primary blocker-hERG docking interfaces within the ion conduction pathway (Figure 4), as follows:

1. The region of the lumen enclosed by the four two-helix bundles of the C-linker residing adjacent to the intra-cellular pore entrance (denoted “C”). L- and Y-shaped blockers likely project moieties (denoted “BC”) into this region, whereas linear blockers may not.
2. The pore lumen spanning between the intra-cellular entrance and intra-cellular face of Tyr652 (denoted “P”). Blockers project a single quasi-linear moiety (denoted “BP”) into this region. The occupied distance along the pore axis varies among blockers.
3. The upper region of the pore, adjacent to the side chains of Tyr652, which we refer to as “Y”). Blockers project the terminal BP group into this region (denoted “BY”), noting that only a subset of blockers terminate in this region (i.e. those that maximally occupy P).

**Figure 3.**
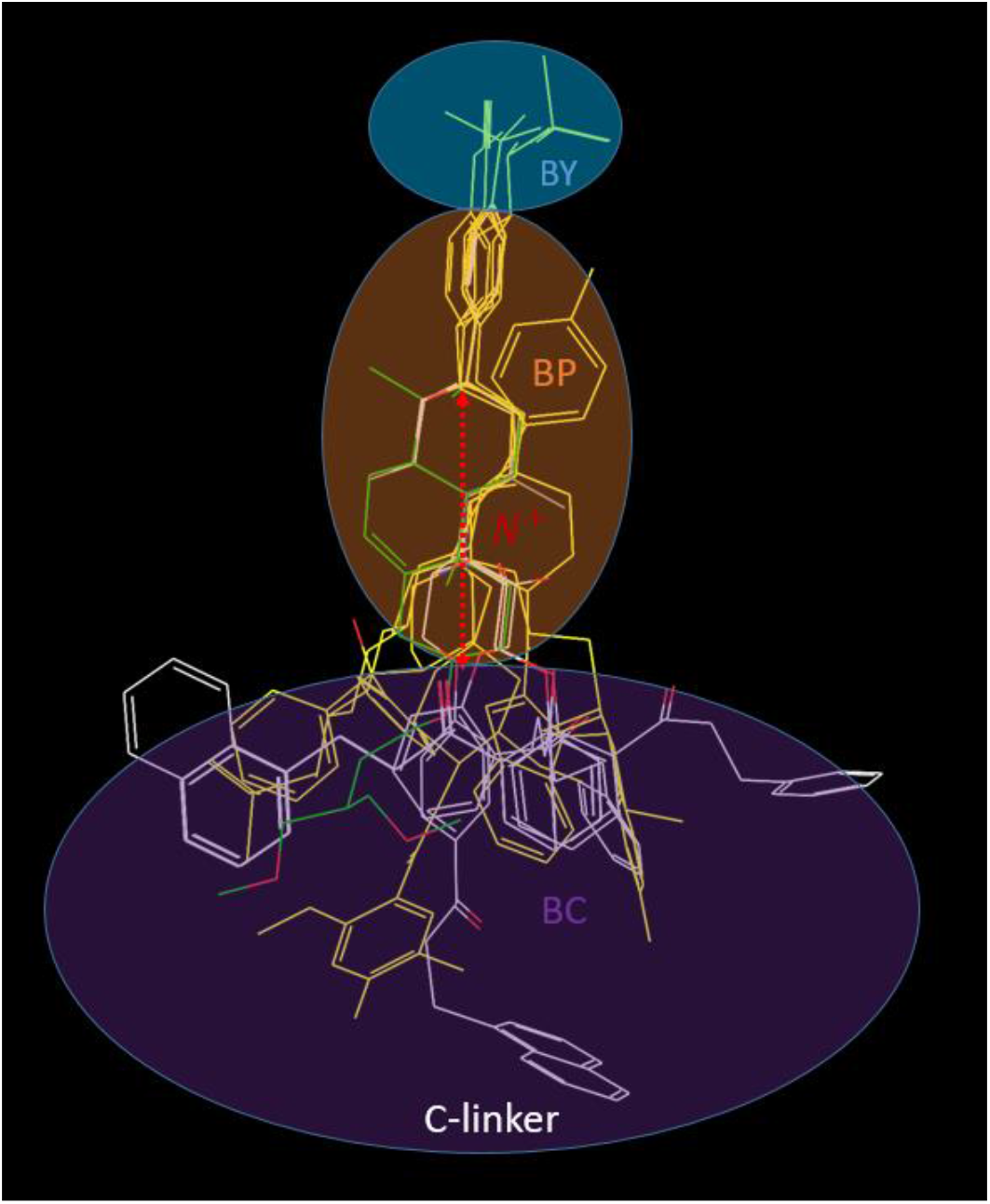
Overlay of the reference set of blockers (see Materials and methods) in the proposed canonical binding mode, which is similar to the CoMFA model reported by Cavalli et al. [32]. BP consists of diverse linear or mildly kinked blocker moieties, typically consisting of one or more hetero-atom containing planar/aromatic or saturated rings, which may be fused (e.g. spiro) or connected by linkers. BC likewise consists of diverse substructures, including butterfly-shaped bisaryl groups.

**Figure 4.**
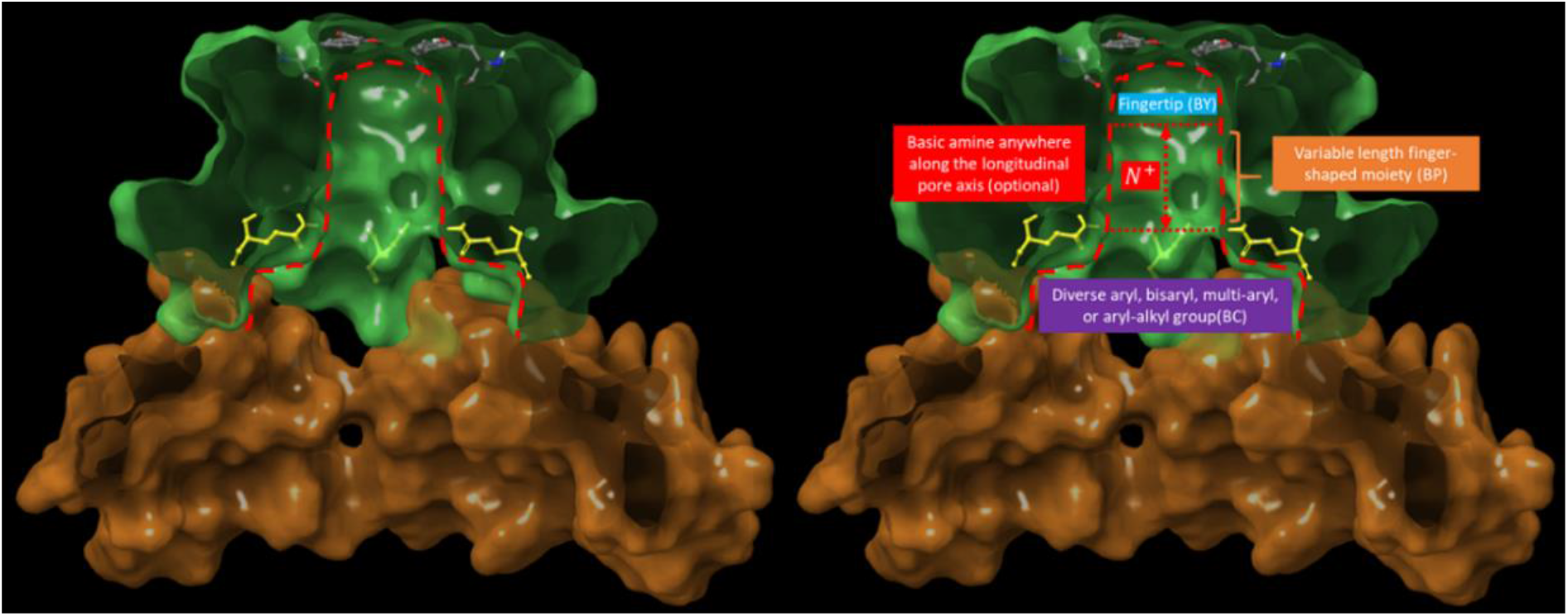
Longitudinal cutaway of the pore and funnel-shaped C-linker cavities (highlighted in green and orange, respectively), showing Tyr652 (comprising the proposed Y docking site) residing at the top of the pore, proximal to the intra-cellular face of the selectivity filter, and Gln664 (highlighted in yellow), which lines the narrowest region of the open pore, and restricts the translocation of BC into the pore. The pore (enclosed within the red dotted outline) is shown without and with annotations for clarity (left and right panels, respectively).

We manually docked terfenadine in the proposed binding mode (Figure 5A) guided by SiteMap site points (the white spheres in Figure 5B).

**Figure 5.**
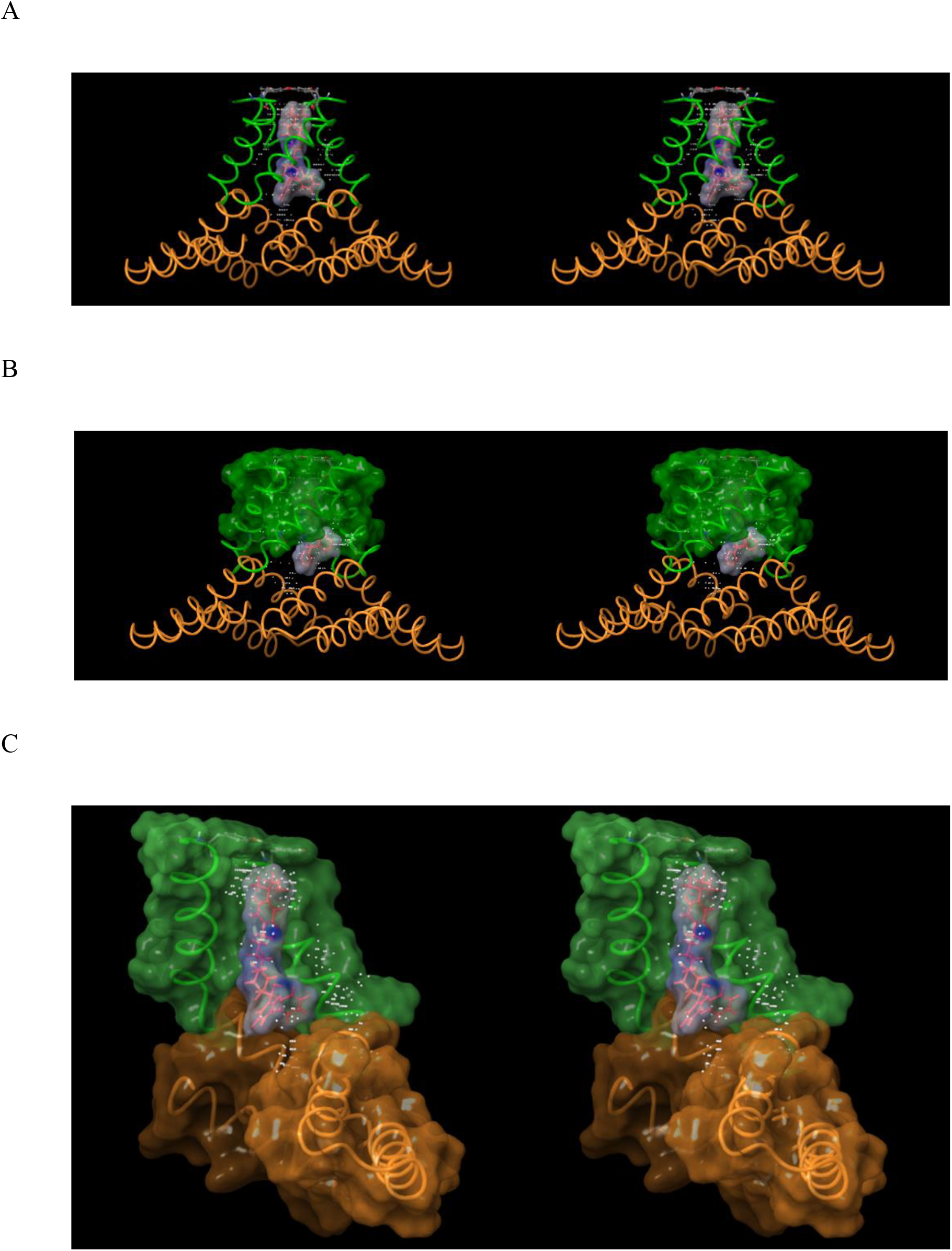
(A) Stereo image of the proposed canonical hERG binding mode (exemplified by terfenadine, shown as a molecular surface), in which blockers straddle between the Y and C regions of the pore (where the C region is comprised of the kink between the S6 and proximal Clinker helices). (B) Same as A, but showing the putative protrusion of the tail region of terfenadine from the pore domain (green surface) into the C-linker enclosed portion of the outer vestibule. (C) Terfenadine was docked <u>manually</u> into clusters of SiteMap site points (white spheres) described in Materials and methods. The butterfly-shaped diphenylmethane tail moiety of terfenadine is complementary in shape to the C-linker helix (noting that butterfly-shaped bisaryl groups are relatively commonplace among hERG blockers [17]).

Blockers necessarily translocate into the pore via the outer vestibule, relegating hERG blockade to a two-step process consisting of:

1. A capture step, in which a single solvated blocker copy diffuses from the cytoplasm into the CNBH domain cavity (which in and of itself is unlikely to block ion conduction into and through the pore).
2. A longitudinal translocation step, in which the captured copy shifts from the CNBHD cavity into the C-linker cavity and pore entrance (Figure 6A-B), while simultaneously:

a. Projecting BP into the open state of P, accompanied by full or partial mutual desolvation of P and BP. Basic groups (when present) reside on BP. BY, the terminal group of BP projects into Y, accompanied by mutual de-solvation of BY and Y. The extent to which BP penetrates into the pore is determined by its length relative to that of P, or the steric size of BY relative to the diameter of the pore entrance.
b. Positioning BC into C, accompanied by full or partial mutual de-solvation of C and BC (<u>noting that BP insertion and BY binding are interdependent processes</u>). Putative H-bond enriched solvation of BC is represented as a red “bumper” in Figure 6C. BC is restricted to the C-linker cavity via steric clashing with side chains at the pore entrance (color-coded yellow in Figure 6), and as such, blockers containing BP moieties shorter than the longitudinal pore length necessarily terminate below the Y docking site.

**Figure 6.**
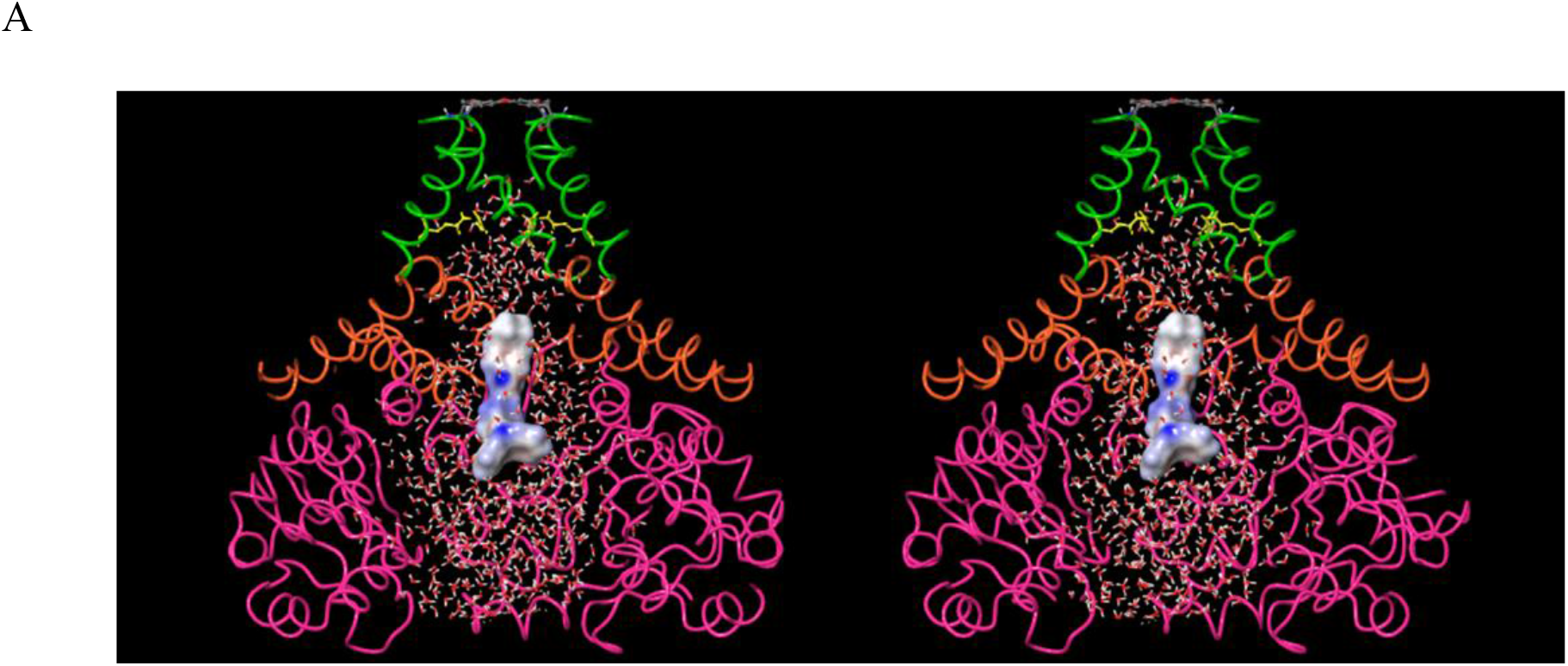
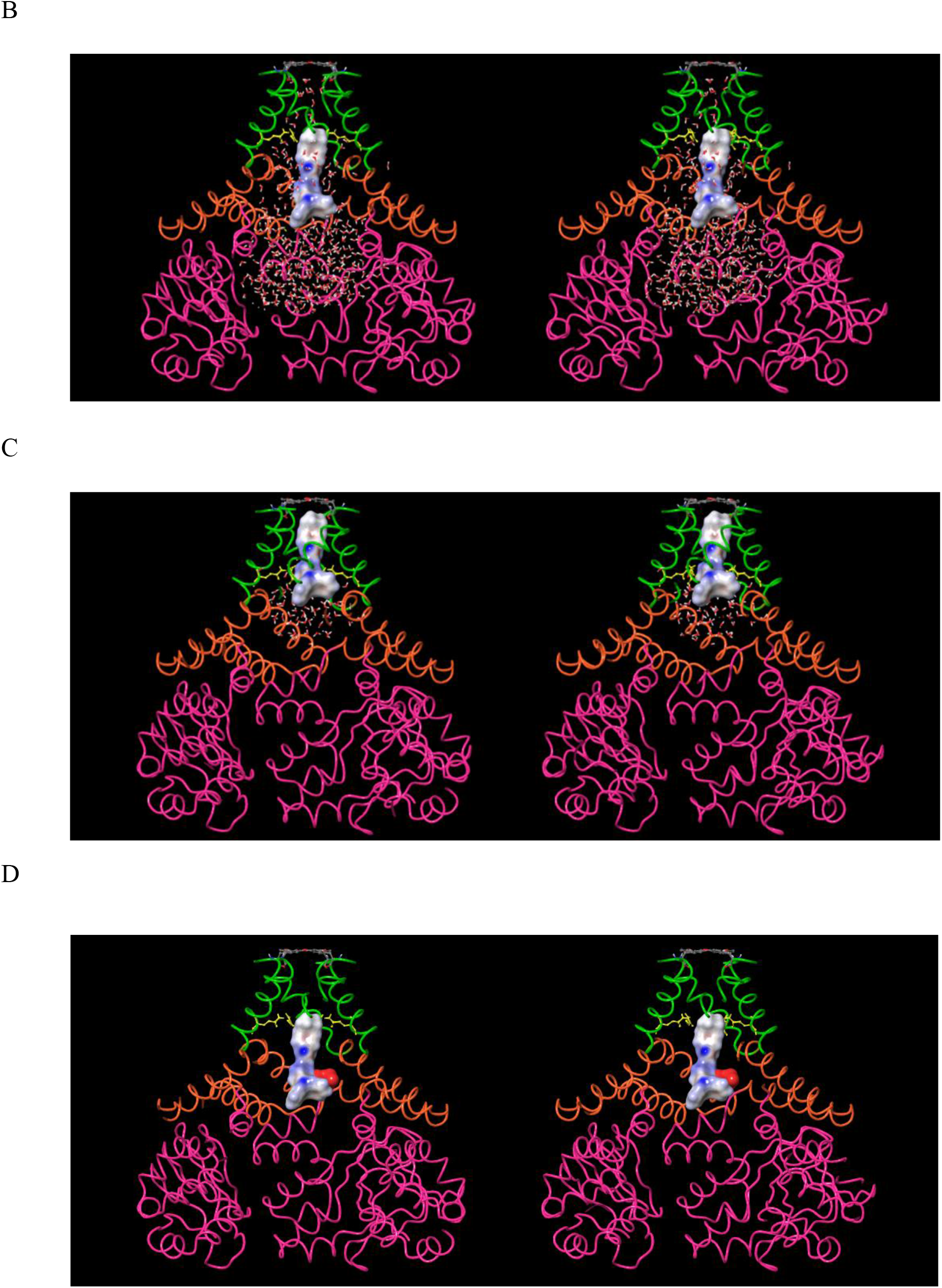
Stereo images of terfenadine manually docked within the cryo-EM hERG structure (PDB code 5VA1 [26]) at various hypothetical stages of binding (and de-solvation), viewed along the pore axis in the intra-(bottom) to extra-cellular (top) direction. (A) Fully solvated terfenadine, manually docked within the CNBH domain region of the outer vestibule prior to putative repositioning of BC, BP, and BY into their corresponding C, P, and Y docking sites. Occupancy of the outer vestibule per se need not block I_Kr_. (B) Terfenadine manually docked with its partially de-solvated BP region penetrating above the pore entrance. (C) Terfenadine manually docked in its hypothetical final binding mode, with the fully de-solvated BC, BP, and BY regions in contact with the C, P, and Y docking sites, respectively. Four such configurations are conceivable due to the 4-fold symmetry of the channel. (D) Cartoon depicting the putative H-bond enriched solvation of BC (red surface). The degree of H-bond enrichment determines the maximum de-solvation free energy cost (the barrier to translocation of BC into C, which behaves like a “bumper” between the two entities in the absence of optimal H-bond replacements).

The total mutual de-solvation costs of C-BC, P-BP, and Y-BY putatively serves as the overall blocker association barrier. The results of our previous WaterMap calculations suggest that P is solvated almost entirely by bulk-like and H-bond depleted water, thereby avoiding disruption of the negative field by ordered H-bond enriched water [17]. The outer vestibule was omitted in our WaterMap calculations, which were performed prior to determination of the cryo-EM structure. H-bond depleted solvation is localized to the non-polar side chains of the pore, and most notably the intra-cellular facing surface of Tyr652 (the Y docking site) located adjacent to the selectivity filter at the distal end of P. We used SiteMap to characterize the solvation within the pore of the cryo-EM structure (data not shown). As expected, the results are consistent with those of our previous WaterMap calculations [17].

### Proposed blocker structure-kinetics relationships

We proposed previously that non-covalent association and dissociation free energy barriers consist principally of H-bond enriched and depleted solvation free energy (relative to the free energy of bulk solvent), respectively [23]. The rate of blocker association depends on the total solvation free energy of H-bond <u>enriched</u> water expelled from the binding interface (i.e. the mutual blocker-channel de-solvation cost). The rate of blocker dissociation depends on the magnitude of the total free energy cost of re-solvating H-bond <u>depleted</u> positions within the dissociated binding interface (i.e. the total blocker and channel re-solvation cost). H-bond <u>enriched</u> solvation incurs zero <u>resolvation</u> cost during <u>dissociation</u>, whereas H-bond <u>depleted</u> solvation incurs zero <u>de-solvation</u> cost during <u>association</u>. In our previous work, we demonstrated that the pore in hERG is solvated almost exclusively by H-bond depleted and bulk-like water [17] corresponding to low de-solvation and high re-solvation costs, respectively. The rate of non-trappable blocker binding, therefore, depends largely on the de-solvation cost of the pore-binding blocker moiety, and the dissociation rate is proportional to the channel-closing rate or blocker dissociation rate, whichever is faster (minimally ~2 s^-1^, which we refer to as the “k_off_ floor” [17]). The rate of pore insertion by trappable blockers is likewise proportional to the de-solvation cost of the pore-binding moiety, whereas the k_off_ floor is ~0.7 s^-1^ [17]. It is therefore apparent that dynamic blocker occupancy is influenced heavily by the rates of channel opening and closing, which is zero in competition binding (where the channels are static), and typically sub-physiological in patch clamp assays. As such, binding measurements performed under non-physiological conditions do not translate reliably to the *in vivo* setting for non-trappable blockers exhibiting k_off_ < the floor <u>and</u> k_on_ < the rate of channel opening. Percent inhibition in all such cases is increasingly overestimated as k_on_ decreases relative to the channel-opening rate (see [17,18]). Blocker binding kinetics can be qualitatively inferred from conventional structure-activity relationships (neglecting channel-gating dynamics), as follows:

1. Significant <u>decrease</u> in the percent inhibition/occupancy of a given blocker analog:

a. k_on_ slowing due to <u>increased</u> blocker <u>de-solvation</u> cost incurred during association via the <u>addition of, or increased polarity of, a polar</u> BC, BP, or BY blocker group (especially BY). Examples of reduced hERG activity putatively due to increased polarity are available in [33] and [34].
b. k_off_ speeding due to <u>decreased</u> blocker <u>re-solvation</u> cost during dissociation via the <u>deletion of, or decreased polarity of, a polar</u> BC, BP, or BY blocker group (especially BY).
c. Slowed k_on_ due to reduced pKa of a basic group (or removal thereof) within the BP moiety.
2. Significant <u>increase</u> in the percent inhibition of a given blocker analog:

a. k_on_ speeding due to <u>decreased blocker de-solvation cost</u> incurred during association via the <u>deletion of, or decreased polarity of, a polar</u> BC, BP, or BY blocker group (especially BY).
b. k_off_ slowing due to <u>increased</u> blocker and/or hERG <u>re-solvation</u> cost during dissociation via the <u>addition of, or decreased polarity of, a non-polar</u> blocker group (especially in BY).
c. k_on_ speeding due to increased pKa of a basic group (or addition thereof) within the BP moiety.

We inferred certain structure-kinetics relationships from the structure-activity relationships of two proprietary in-house datasets based on the aforementioned principles. A large activity cliff in dataset 1 is attributable to increased de-solvation cost of the pyridazyl versus pyridyl moieties predicted to bind in site C (Table 4) and the 1-versus 2-pyridyl combined with Cl versus F (Table 5). The structure-property relationships underlying the solvation differences among these groups is unobvious. A large activity cliff in dataset 2 can be attributed to increased de-solvation cost of the oxadiazole versus sulfadiazole and oxapyrrole groups predicted to project into a cluster of H-bond depleted solvation in site Y (Table 6), which putatively manifests as slowed k_on_ due to the loss of H-bonds of polar group solvation transferred to bulk solvent. We note that key local solvation effects may be masked in global changes in <u>scalar</u> logP and solubility. Non-trappable blocker-induced expulsion of this solvation is expected to slow k_off_, minimally to the rate of channel closing (below which channel closing becomes rate-determining). k_off_ slowing in the case of shorter BP groups falling short of the Y site depends on the expulsion of H-bond depleted solvation from the C region (equating to an enthalpic <u>re-solvation</u> cost during dissociation), together with the entropic de-solvation contribution.

**Table 3.**
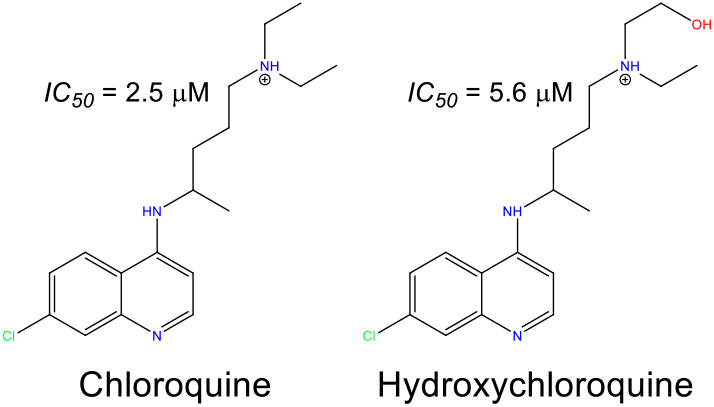
Candidate COVID-19 therapies known to block hERG [28,29] (noting that the HCQ IC_50_ was estimated from the reported percent inhibition (35%) at 3 μM using the Hill equation).

**Table 4.**
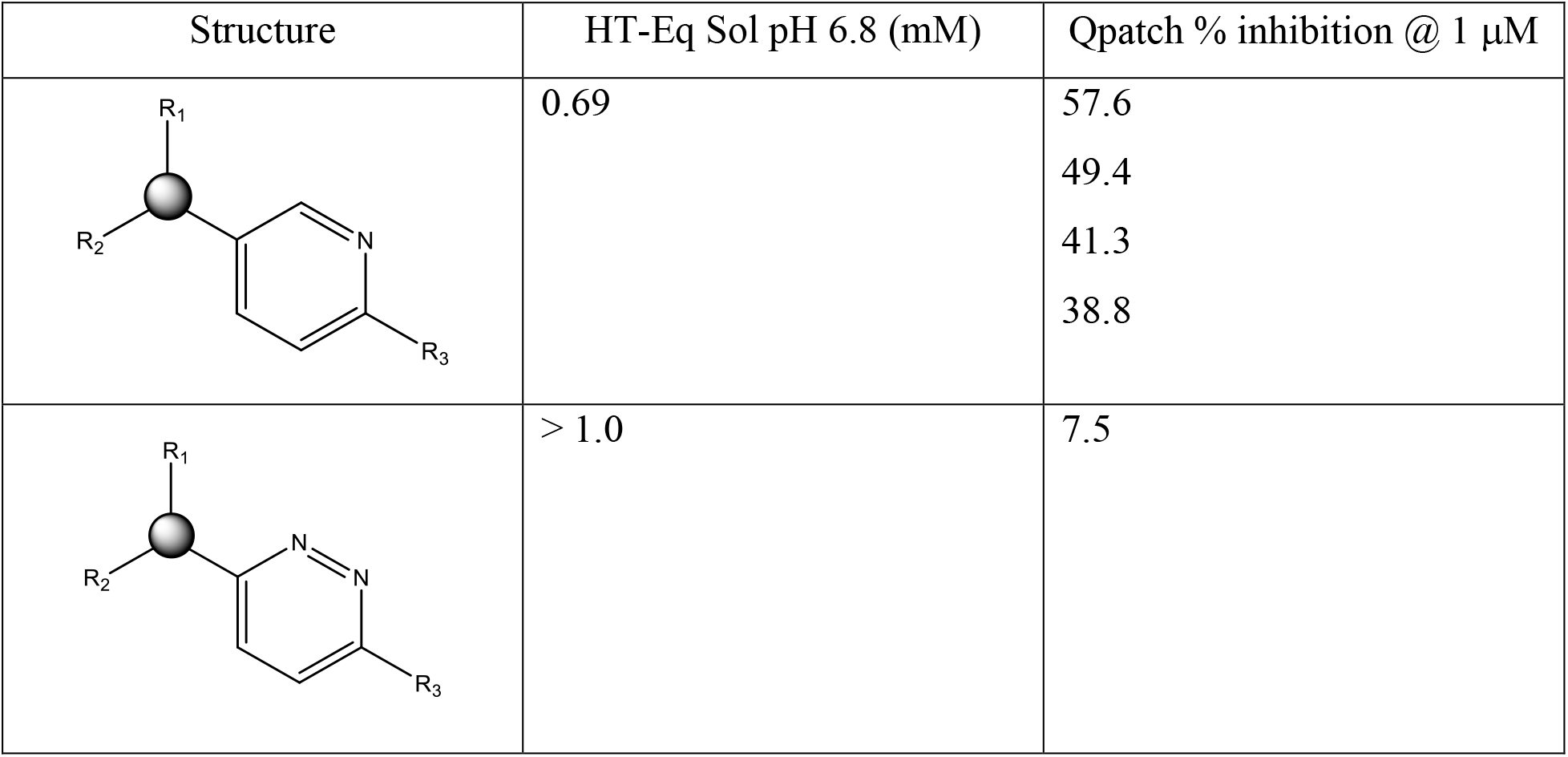
Representative activity cliffs for a proprietary hERG blocker series differing solely in the six-ring substructures (residing within the putative BC feature set). The large decrease in percent inhibition @ 1 μM for structure 2 compared with structure 1 is attributable to increased polarity and solubility/de-solvation cost of the pyridazyl versus the pyridyl group (consistent with the lack of hERG-contributed H-bond replacements for the water solvating this moiety in the unbound state).

| Structure | HT-Eq Sol pH 6.8 (mM) | Qpatch % inhibition @ 1 $\mu\text{M}$ |
| --- | --- | --- |
|  | 0.69 | 57.6<br>49.4<br>41.3<br>38.8 |
|  | > 1.0 | 7.5 |

**Table 5.**
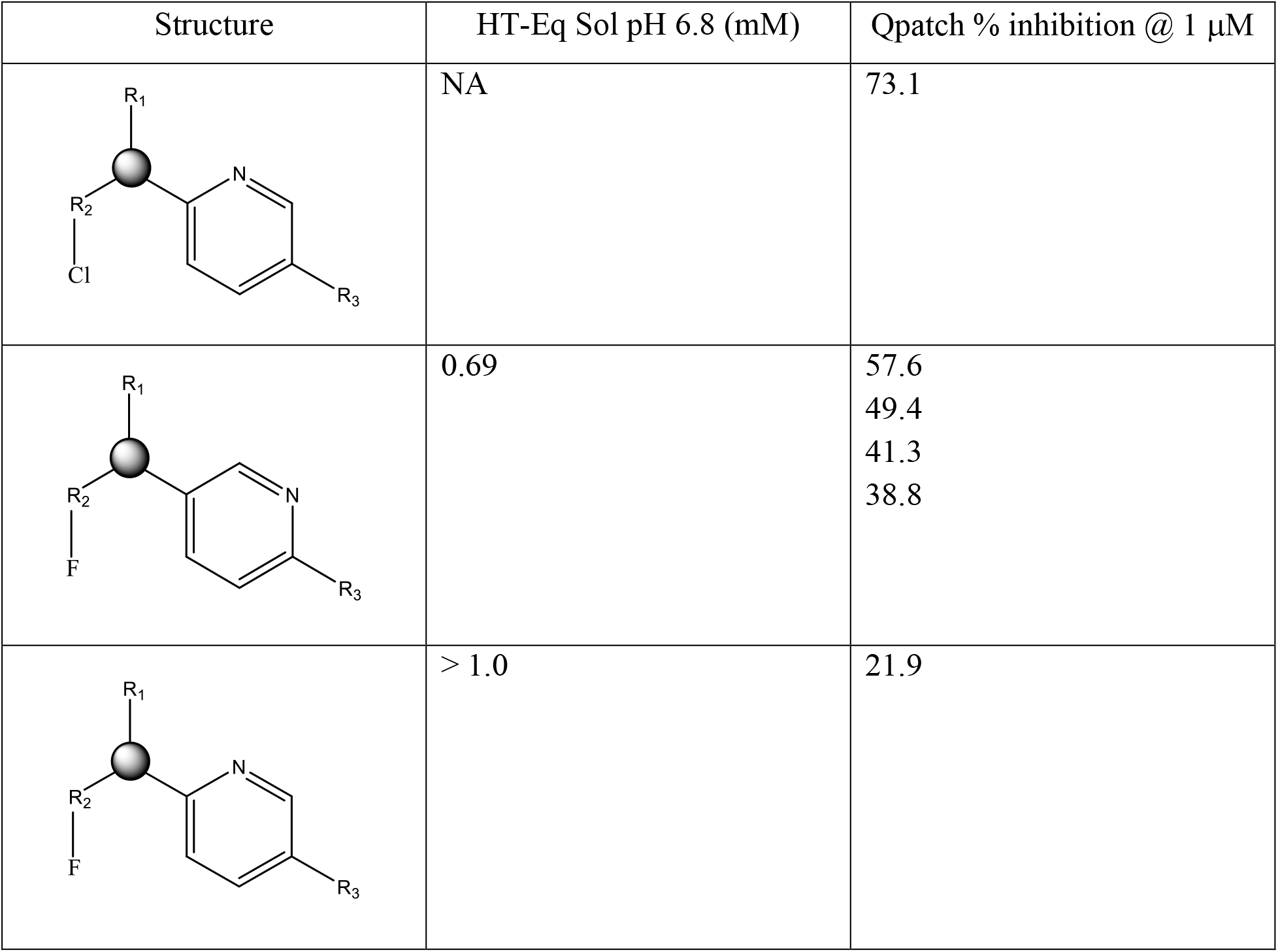
Representative activity cliffs for the proprietary hERG blocker series 1 (see text) comprised of R1 (corresponding to BP) and R2/R3 (corresponding to BC). The large decrease in percent inhibition @ 1 μM for structure 3 compared with structures 1 and 2 is putatively attributable to the greater re-solvation cost of Cl versus F, together with increased polarity and solubility/de-solvation cost of the ortho-versus meta-pyridyl nitrogen, (consistent with the lack of hERG-contributed H-bond replacements for the water solvating this moiety in the unbound state).

| Structure | HT-Eq Sol pH 6.8 (mM) | Qpatch % inhibition @ 1 μM |
| --- | --- | --- |
|  | NA | 73.1 |
|  | 0.69 | 57.6<br>49.4<br>41.3<br>38.8 |
|  | > 1.0 | 21.9 |

**Table 6.**
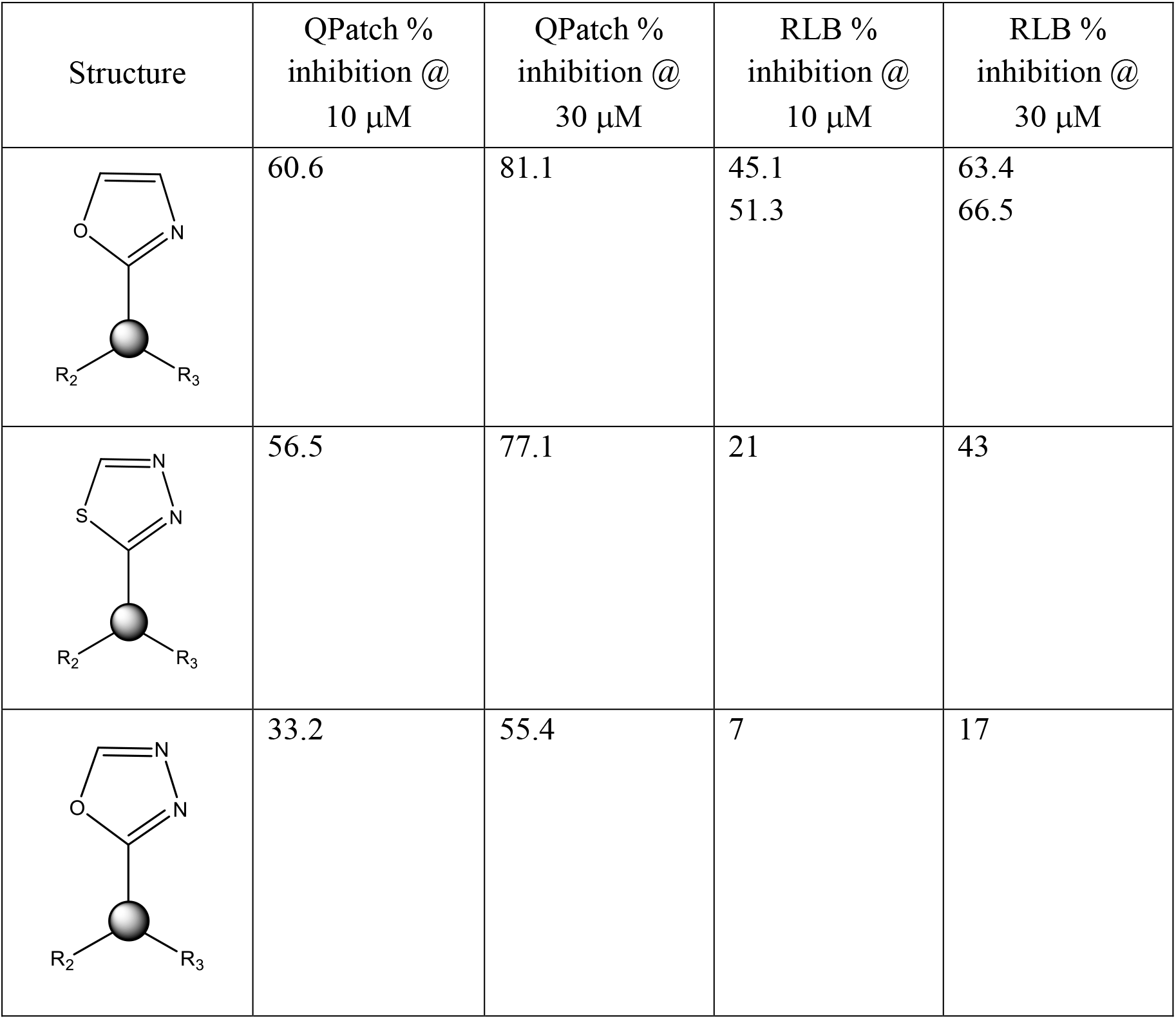
Representative activity cliffs for the proprietary hERG blocker series 2 (see text) comprised of R1 (corresponding to BP) and R2/R3 (corresponding to BC). The data suggests that hERG occupancy is disrupted by the projection of a highly polar group (oxadiazole) versus a less polar group (sulfadiazole or oxapyrrole) into the putative localized region of H-bond depleted solvation at site Y.

| Structure | QPatch %<br>inhibition @<br>10 μM | QPatch %<br>inhibition @<br>30 μM | RLB %<br>inhibition @<br>10 μM | RLB %<br>inhibition @<br>30 μM |
| --- | --- | --- | --- | --- |
|  | 60.6 | 81.1 | 45.1<br>51.3 | 63.4<br>66.5 |
|  | 56.5 | 77.1 | 21 | 43 |
|  | 33.2 | 55.4 | 7 | 17 |

### Proposed blocker structure-trappability relationships

The concept of blocker trappability was proposed by Starmer et al. [35] and Stork et al. [36] (referred to by Armstrong et al. [37] and Mitcheson et al. [38] as the “foot-in-the-door” model). Non-trappable and trappable blockers are differentiable in patch clamp experiments on the basis of channel gating frequency-dependent versus independent inhibition, respectively. Higher frequency stimulation results in proportionately greater fractional open channel time, and therefore, increased binding site accessibility to non-trappable blockers that build and decay during each gating cycle (noting that trappable blockers accumulate to their steady-state occupancy regardless of the open channel time, merely accumulating faster with increasing open time). Windisch et al. studied structure-trappability relationships for a series of analogs around the known trappable blocker, propafenone [25]. We set about to explain this relationship by overlaying and comparing the predicted conformational properties of these analogs vis-à-vis our proposed canonical drain-plug model (see Materials and methods for a description of the conformational analysis approach). The results unambiguously demonstrate that trappable versus non-trappable propafenone analogs exhibit planar versus non-planar conformations, respectively, at the position occupied by the uppermost phenyl ring in Figure 7. Furthermore, the trappability-determining region in the full set of overlaid blockers maps to a specific region of the pore (Figure 8) that we hypothesize constricts in the closed channel state and sterically clashes with non-planar blocker moieties that are otherwise compatible in the open state. We speculate that the putative constriction zone is comprised of Phe656 side chains that rearrange from peripheral positions observed in the cryo-EM structure to more central positions during channel closing. This hypothesis is supported by a model that we generated of the closed form of the channel (results to be reported in a future work).

**Figure 7.**
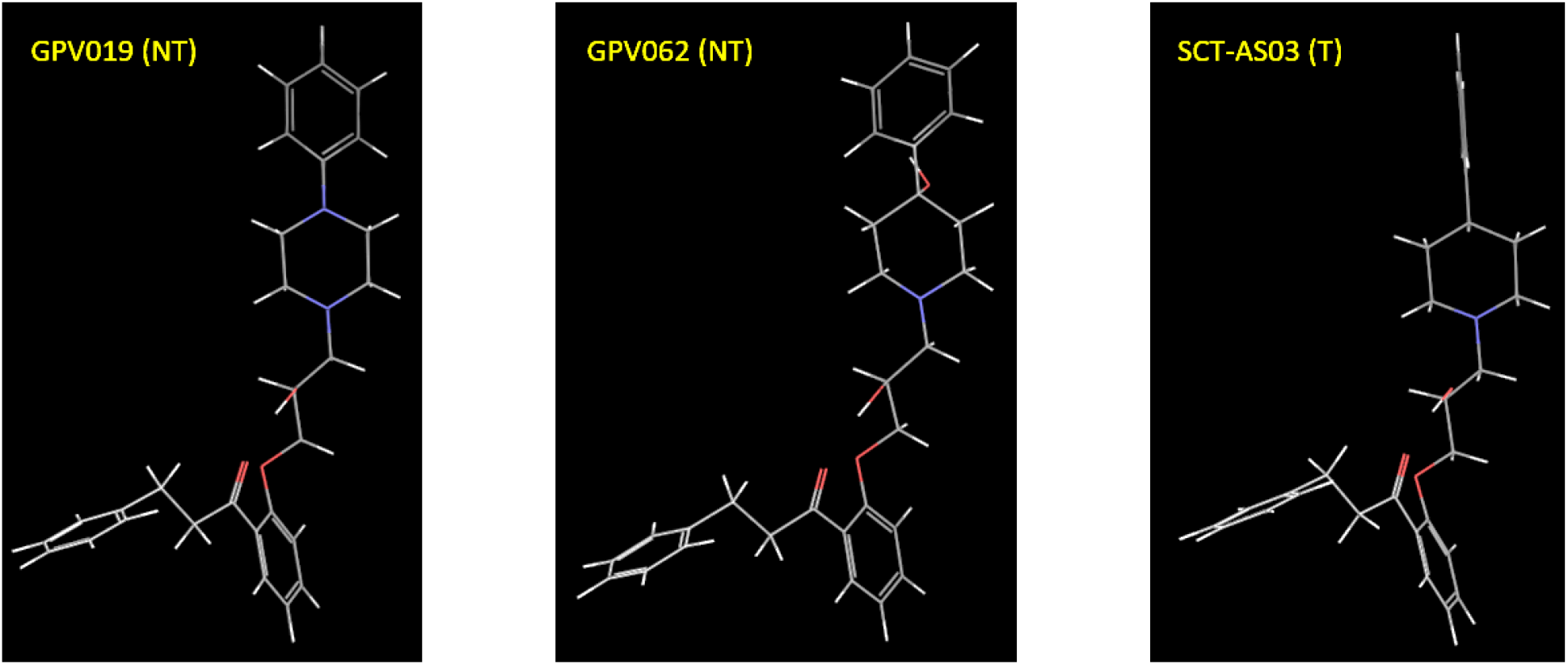
Unambiguous conformational differences are predicted between measured trappable (T) and non-trappable (NT) propafenone analogs. In particular, trappable blockers such as SCT-AS03 are highly planar within the proposed “constriction zone,” whereas non-trappables, such as GPV019 and GPV062, are not.

**Figure 8.**
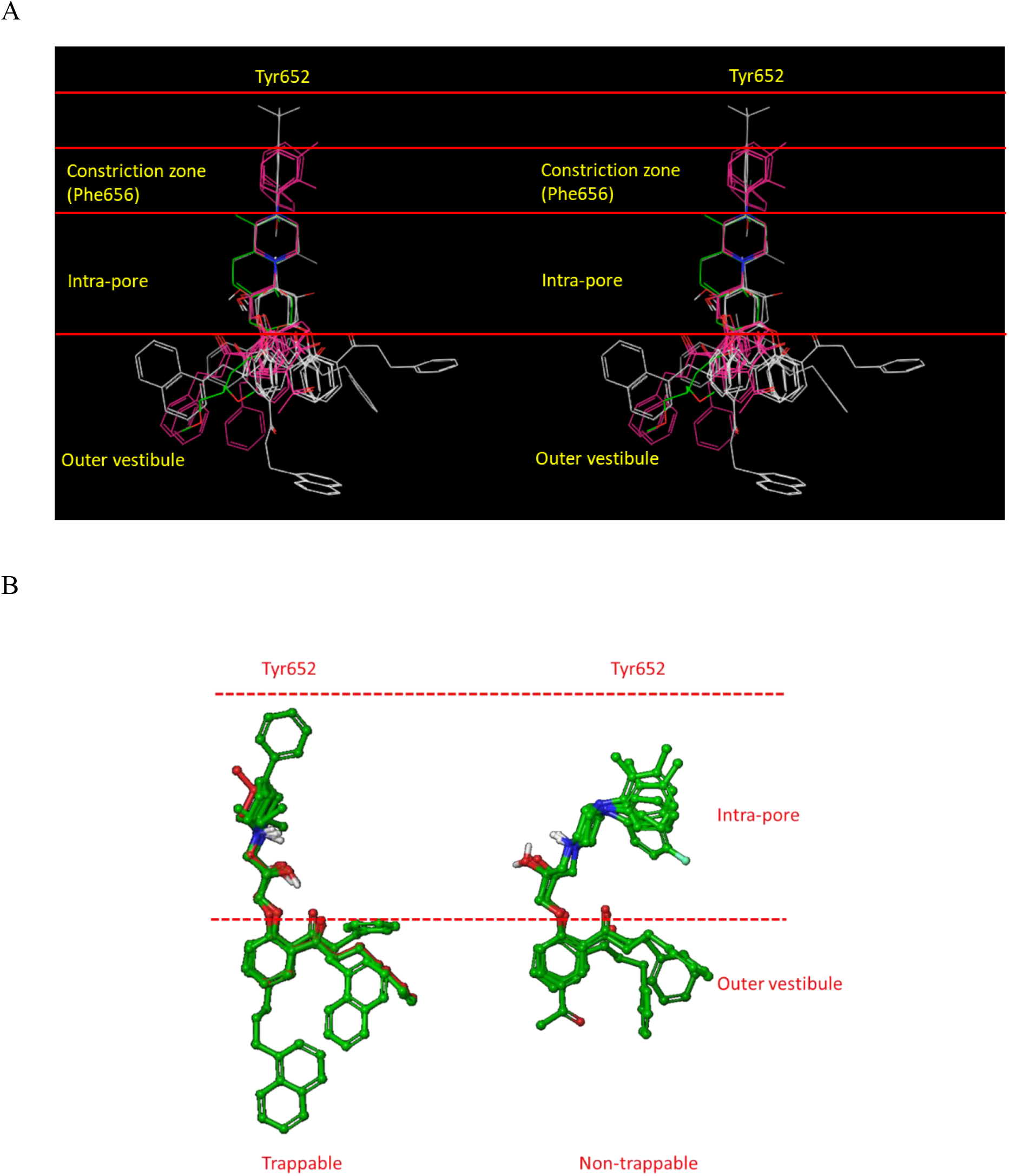
Our blocker overlay model (Figure 3) mapped to the various zones of the channel, including the putative trappability-determining constriction zone that forms during channel closing, from which non-trappable blockers are sterically excluded. (A) Stereo view of the blocker overlay. All of the trappable propafenone analogs either terminate prior to this zone or project a planar group into it (magenta). (B) Zoomed-in view of the putative blocker constriction zone, demonstrating the non-planar conformations of non-trappable blockers in this region.

In summary, blocker binding is driven largely by:

1. Concurrent steric shape complementarity to the C (L- or Y-shaped blocker moieties), P (quasi-linear blocker moieties), and Y regions of the channel.
2. Principally <u>blocker</u> de-solvation costs during translocation from the CNBHD cavity into the C, P, and Y (optionally) regions of the channel.
3. Principally, <u>channel</u> re-solvation costs at the Y and C sites incurred during blocker-hERG dissociation.
4. Electrostatic interactions with basic blockers that are capable of projecting their charged group along the longitudinal axis of P (Figures 3 and 4, right).

Trappable blockers either terminate within P prior to the putative constriction zone formed by Phe652 side chains, or project a planar group into the constriction zone. Trappable blockers accumulate occupancy over multiple channel gating cycles (i.e. heartbeats), peaking at the <u>intracellular</u> free C_max_, whereas non-trappable blocker occupancy builds and decays within the ventricular hERG population during each cycle. The maximum dynamic occupancy (corresponding to k_off_/k_on_) <u>across all cycles</u> within a dosing period peaks at the free intra-cellular C_max_ for both trappable and non-trappable blockers. Trappable blocker occupancy is agnostic to gating frequency (i.e. on-rate merely governs the rate of buildup to the maximum occupancy) whereas non-trappable blocker occupancy depends on k_on_ and k_off_ relative to the channel opening and closing rates, respectively.

### Possible deficiencies of the status quo hERG safety assessment protocol

The Redfern SI (equation 1) was derived from a comparison of measured hERG IC_50_, therapeutic C_max_, and maximum observed C_max_ for 100 marketed drugs classified according to TdP propensity: withdrawn drugs versus multiple reported cases, versus isolated cases, versus no reported cases, versus anti-arrhythmic hERG blocking drugs. We have identified the following potential deficiencies in the derivation and application of this metric:

#### Deficiency 1

Substitution of equation 1 into the Hill equation yields a maximum safe hERG blocker occupancy (γ_1_) = 3% [17] (by all accounts, a highly stringent safety margin in the absence of underlying cardiovascular disease):

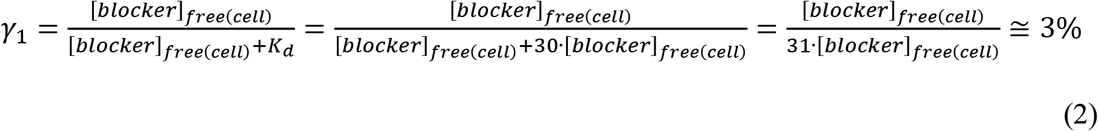

However, it is apparent that the TFPC_max_ equates to the therapeutic <u>total intra-cellular</u> C_max_ (hereinafter referred to as TTIC_max_) (Figure 9). Substitution of [blocker]_free(cell)_ by [blocker]_total(cell)_ - [blocker]_bound(cell)_ into equation 3 results in an underdetermined equation in the absence of [blocker]_bound(cell)_ information:

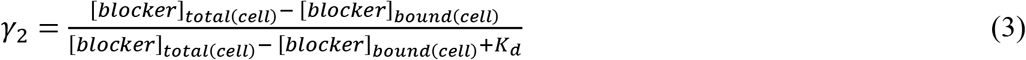

**Figure 9.**
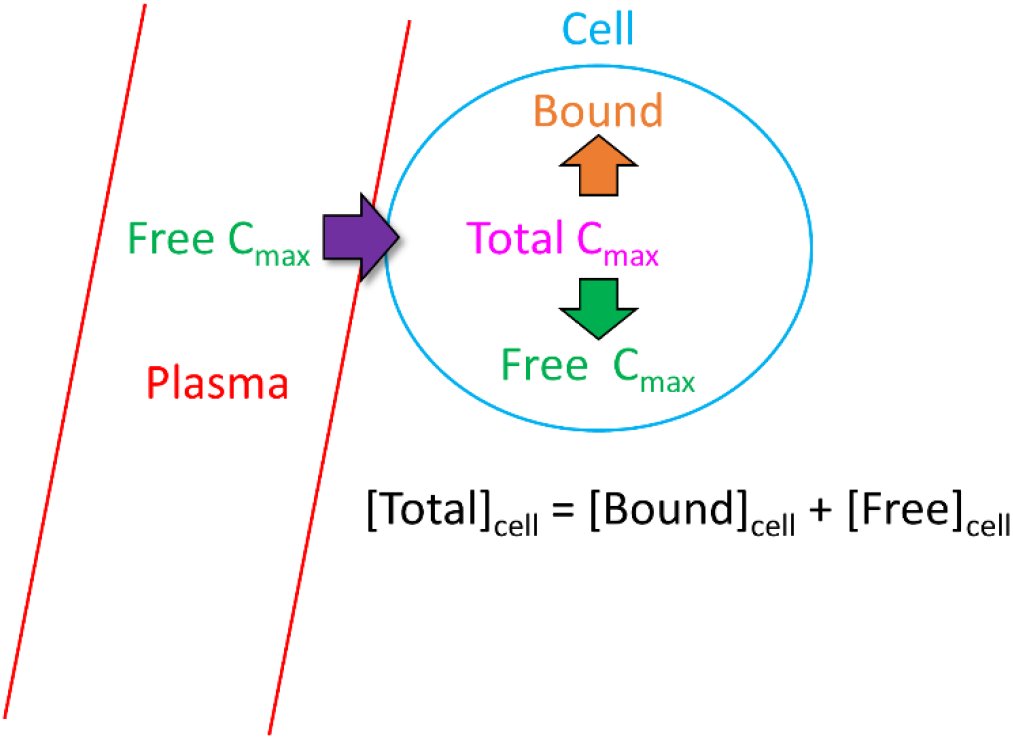
TFPC_max_ equates to TTIC_max_, such that the Hill equation, which is based on free concentration, is unsolvable (see text).

However, the ratio of γ_2_ to γ_1_ is necessarily less than 1 because [blocker]_total(cell)_ in equation 3 is always greater than [blocker]_free(cell)_:

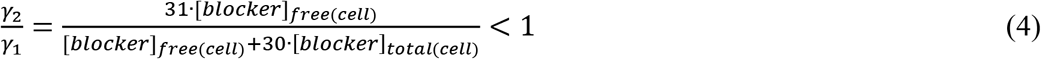

Considering [blocker]_total(cell)_ = 5 · [blocker]_free(cell)_ for example leads to:

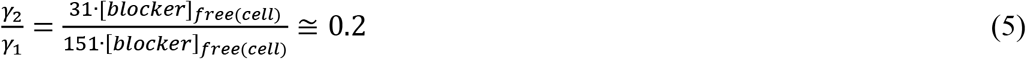

Viewed in this light, the Redfern SI equates to approximately zero safe hERG occupancy at the TFPC_max_ (= TTIC_max_), which buffers exposure and occupancy escalation between the TFPC_max_ and arrhythmic FPC_max_ due to DDI, overdose, or other factors. A wide safety margin of this nature is arguably justified given the life and death implications of drug-induced TdP (exacerbated in the presence of pre-existing cardiac impairment), although possibly tempered by questionable assumptions in the Redfern SI derivation.

#### Deficiency 2

Both the TFIC_max_ (which correlates with the TFPC_max_) and hERG safety margin necessarily translate to a maximum FIC_max_ (at both therapeutic and escalated exposures due to DDI or overdose) far below our predicted arrhythmic FIC_max_ (AFIC_max_) in the undiseased heart. The hERG IC_50_/30 criterion derived by Redfern et al. [39] is implicitly weighted toward the most potent TdP-inducing drugs in the dataset, which correctly accounts for the narrow exposure window between the maximum observed FIC_max_ and AFIC_max_. However, this relationship breaks down with increasing hERG IC_50_ because the <u>absolute</u> exposure window (i.e. Δ(exposure) = arrhythmic FIC_max_ – hERG IC_50_/30) matters, rather than the ratio. In our previous work, we used the O’Hara-Rudy model to predict the cellular arrhythmic threshold of hERG inhibition relative to the open and total hERG populations in otherwise normal midmyocytes (M cells), which varies with cell type (M cells being the most sensitive by far) and blocker trappability [17]. Here, we use a conservative estimate of ~50% occupancy of the open hERG population (translating to ~50% reduction of I_Kr_ at free intra-cellular exposure ≈ hERG IC_50_) for reference, noting that we are currently re-exploring hERG blocker dose-response behavior (results to be reported elsewhere). The Redfern SI can thus be roughly expressed as: Δ(exposure) = hERG IC_50_ -hERG IC_50_/30, which is exemplified in Figure 10 for hERG IC_50_ = 1 nM, 1 μM, and 10 μM. It is apparent from these examples that the upper safe TFPC_max_ in otherwise normal M cells is greatly underestimated for μM blockers by the ratio-based one-size-fits-all Redfern SI compared with absolute Δ(exposure).

**Figure 10.**
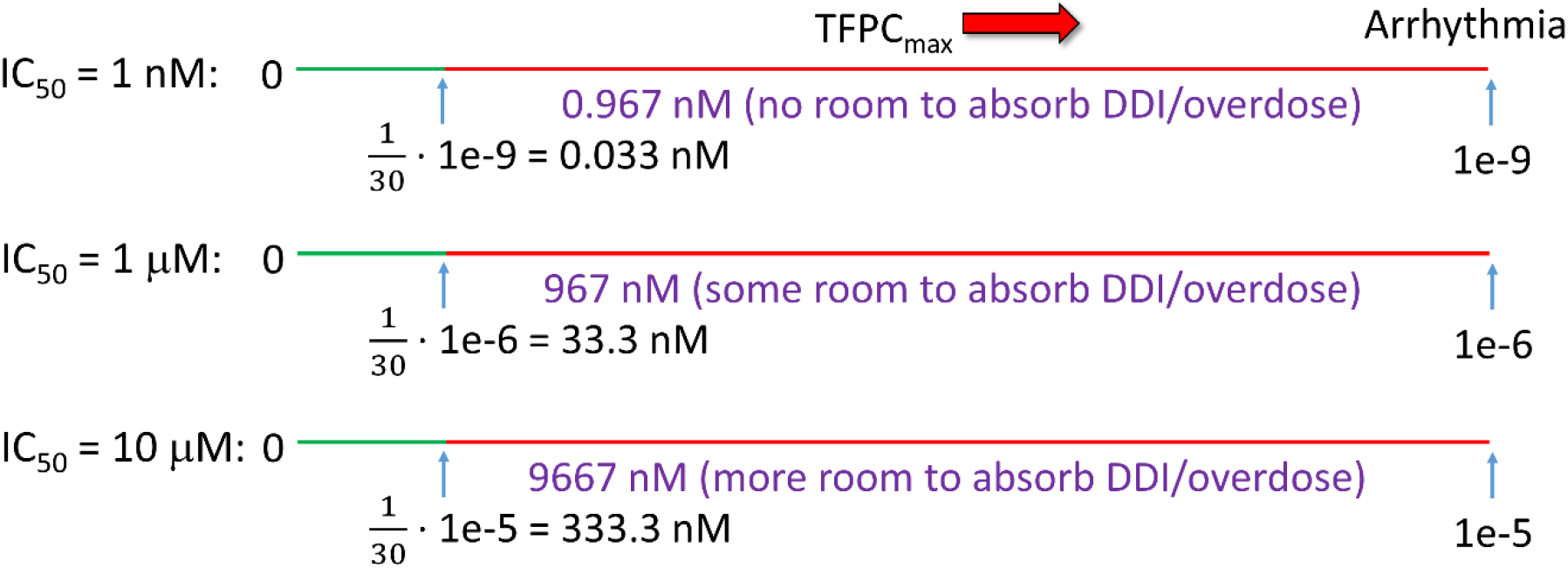
The Redfern et al. derivation is tacitly based on the assumption that the safety margin consists of the exposure range between the TFPC_max_ and the maximum FPC_max_ due to DDI, overdose, or other factors, equating to 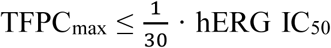. For the present purpose, we assume that arrhythmia (the primary origin of organ-level TdP) occurs at approximately 50% inhibition of the open channel population in otherwise normal M cells (corresponding to the apparent IC_50_), and possibly much lower in the diseased heart, translating to an absolute Δ(exposure)-based Redfern SI = hERG IC_50_ – hERG IC_50_/30.

#### Deficiency 3

Blocker occupancy (proportional to percent inhibition) is channel state- and time-dependent, and therefore out of the scope of equation 3:

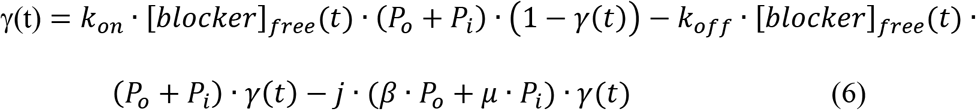

where *[blocker]_free_(t)* is the time-dependent free intra-cellular blocker concentration, *P_o_ and P_i_* are the open and inactive state probabilities, *β* is the channel opening rate constant, and *μ* is the channel inactivation rate constant [17]. Non-trappable blocker occupancy at a given *[blocker]_free_* is overestimated by equation 3 when k_on_ is less than the channel opening rate and *[blocker]_free_* is below the saturating level. Trappable blocker occupancy is described by equation 3 when k_off_ ≥ k_off_ floor (which cannot be ascertained from IC_50_ measurements alone). The *in vivo*-relevant percent inhibition for both trappable and non-trappable blockers is ideally measured via manual patch clamp experiments performed at the physiological gating frequency. Given the fast rate of channel opening (the peak amplitude is reached in a few ms), the k_on_ requirement for high percent inhibition among non-trappable blockers is necessarily fast [17]. Zu et al. measured the rate constants for several hERG blockers (including some listed in Table 1) <u>in static channels</u> via a radio-ligand displacement approach [40]. Using cisapride (a known non-trappable blocker) as a benchmark, we hypothesize that all of the measured rate constants in the authors’ dataset are up to 1,000-fold slower relative to physiological conditions (noting that k_on_ and k_off_ were deconvoluted from the measured kobs = k_on_ · [blocker]_free_ + k_off_ and the IC_50_).

#### Deficiency 4

Status quo hERG mitigation and preclinical safety assessment is typically guided by high throughput *in vitro* IC_50_ data and animal PK data. hERG mitigation depends on guidance from accurate data capable of unambiguously resolving the true SAR [41], which is difficult to acquire via high throughput *in vitro* measurements (Figure 11) (noting the often flat hERG free energy landscape across primary target potency-sparing chemical series). Mitigation is aimed minimally at satisfaction of the Redfern SI, and maximally at total abrogation of hERG activity (i.e. IC_50_ > the detectability limit) among clinical drug candidates. However, total hERG mitigation combined with optimal primary target potency and other requirements, is rarely achieved in practice, relegating high uncertainty to both terms of the Redfern IC_50_/exposure ratio. Uncertainty on top of uncertainty is especially problematic for safety assessment of clinical candidates in advance of *in vivo* cardiotoxicity evaluation.

**Figure 11.**
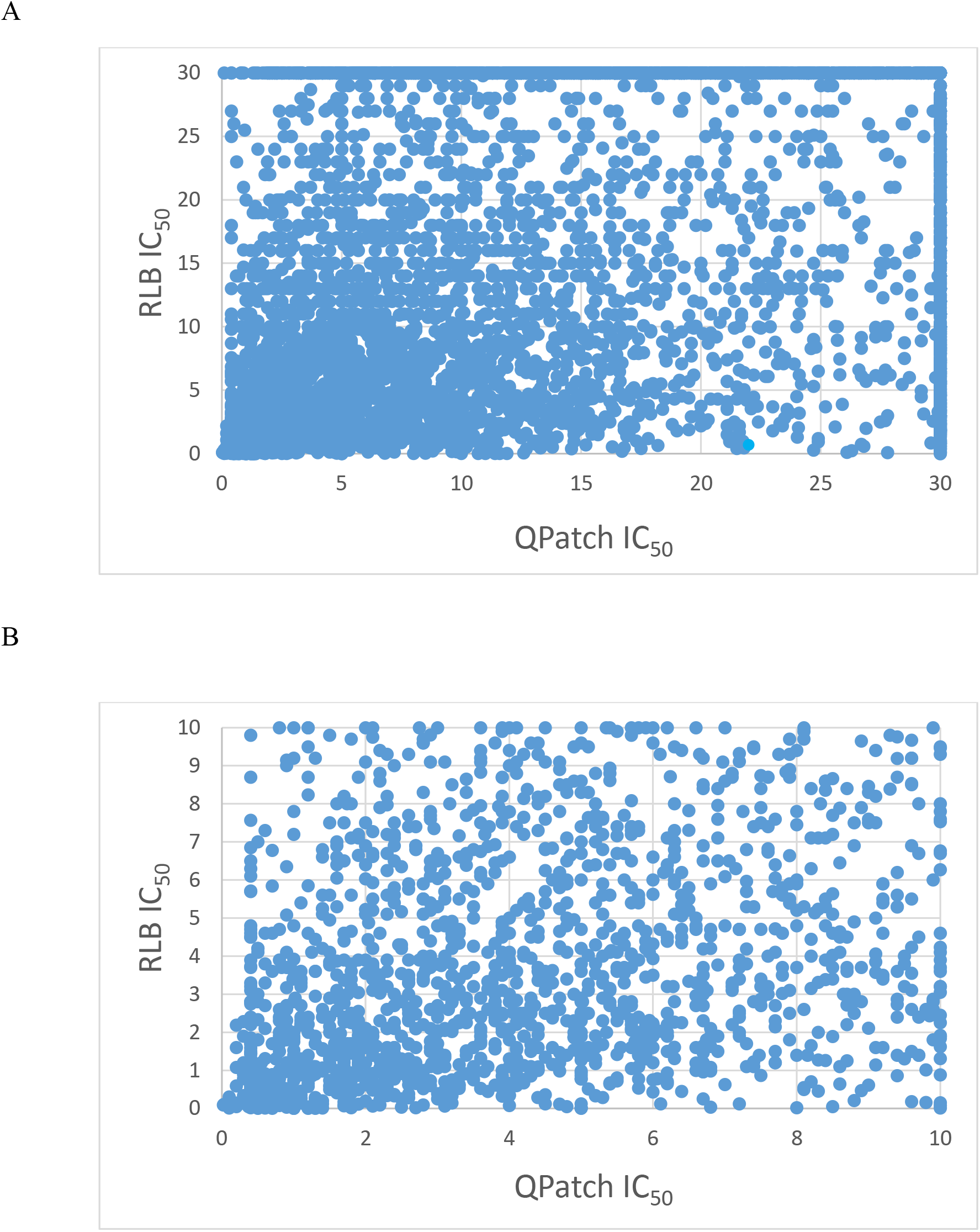
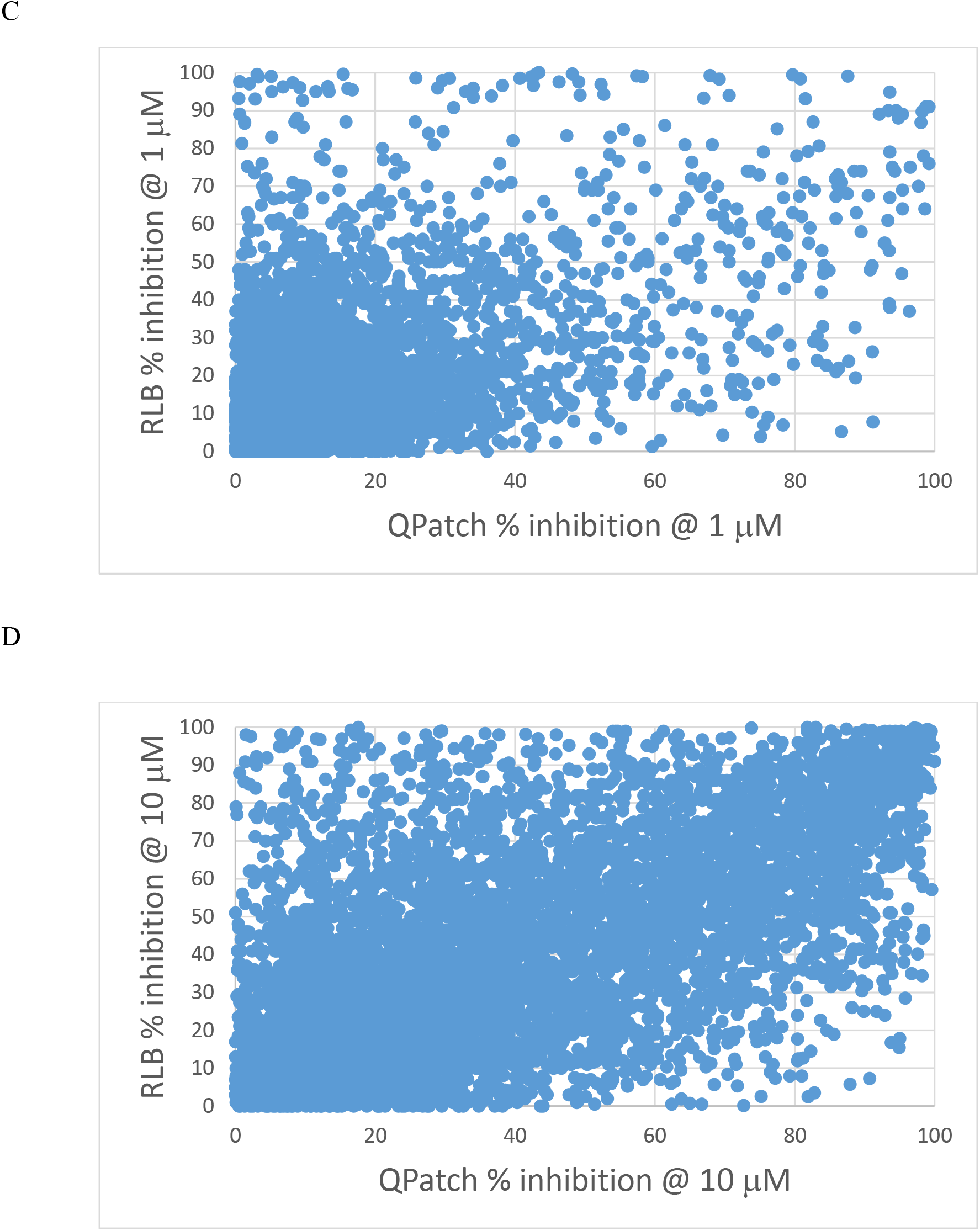
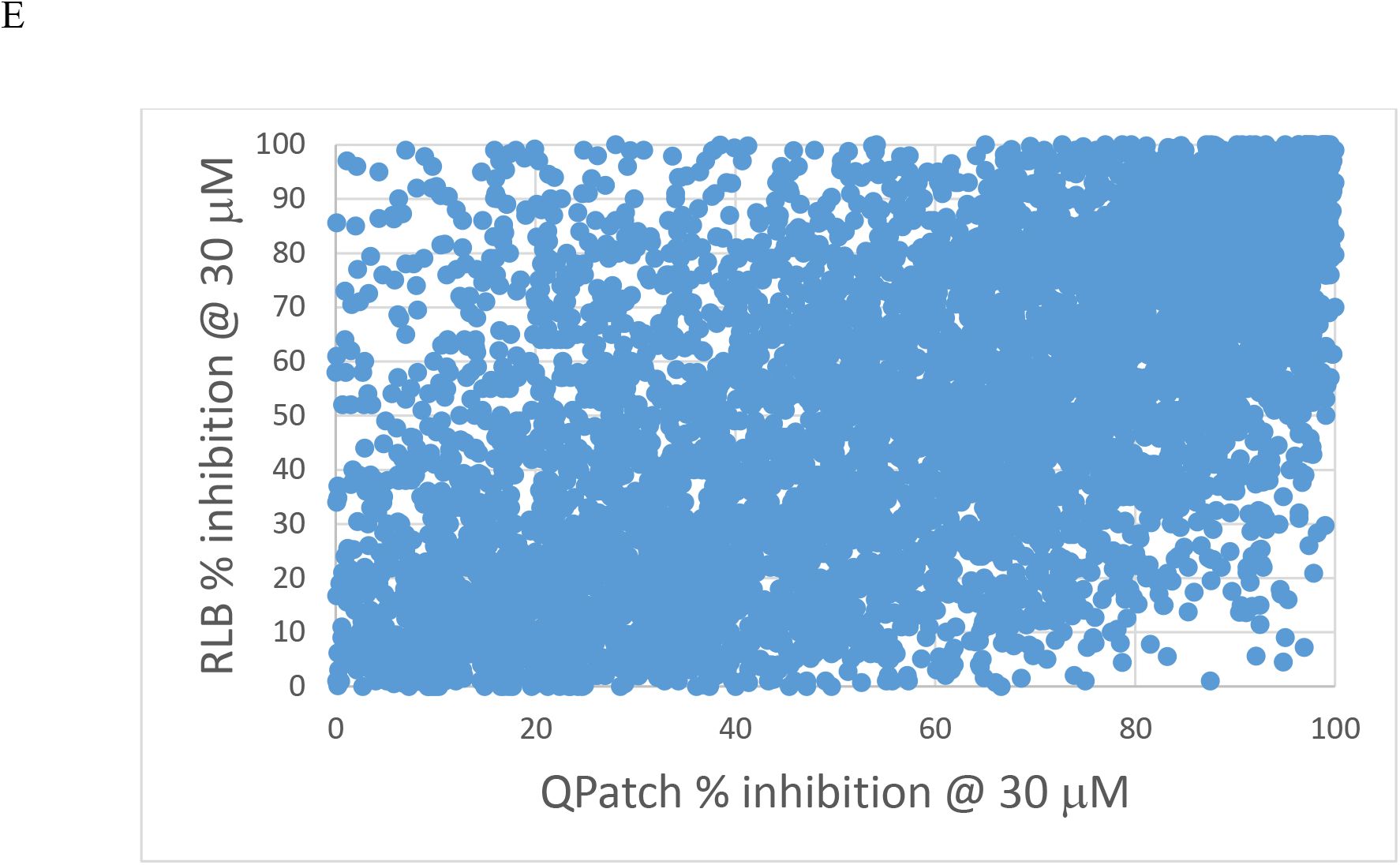
Plots of measured high throughput QPatch versus radio-ligand binding (RLB) assays for 7,231 internal proprietary compounds. (A) IC_50_. (B) IC_50_ zoomed to the 0 to 10 μM range. (C) Percent inhibition @ 1 μM. (D) Percent inhibition @ 10 μM (E) Percent inhibition @ 30 μM. The lack of agreement between the two assays at all percent inhibition levels is apparent. The accuracy/resolution of this data may be insufficient to support SAR analysis needed to guide hERG mitigation, as well as QSAR and machine learning models.

### Proposed clinically-relevant hERG safety assessment and mitigation approaches

As addressed above, pro-arrhythmic risk at exposures ≥ the TFPC_max_ in humans is putatively overestimated by the *in vitro* IC_50_-based Redfern SI in patients with uncompromised cardiac function (a key assumption in our simulations). Uncertainty in the hERG safety of late stage lead compounds exhibiting residual activity is often further complicated by the lack of confirmed quantitative hERG data (which may extend to clinical candidates in some cases). Instead, we propose the following approach:

1. Minimize TFPC_max_ via kinetically tuned <u>drug-target</u> binding, defined as the optimization of k_on_ to the rate of binding site buildup (explained in detail in [17] and Selvaggio et al. [18]).
2. Test for blocker trappability based on the structural criteria outlined above (Figures 7–8), together with frequency independent percent inhibition using an appropriate patch clamp protocol [36].
3. Perform trappable ➔ non-trappable chemical transformations wherever possible based on the structural criteria outlined above (noting that non-trappable blocker occupancy does not accumulate over time).
4. SAR and safety assessment should be based on <u>percent inhibition</u>, rather than IC_50_ data (noting that error may be amplified in best-fit dose-response curves, secondly, that percent inhibition is a more direct measure of blocker-hERG occupancy at concentrations of interest, and thirdly, fractional hERG occupancy *in vivo* cannot be predicted from IC_50_ in the absence of TFIC_max_). Sufficient accuracy is needed to confirm critical SAR and inform safety assessment. Data accuracy can be gauged as follows:

a. Similar observed trends between orthogonal assays (e.g. patch clamp and radioligand binding).
b. Convergence of replicate runs in each assay.
c. Self-consistency of hERG SAR (the existence of identifiable hERG structureactivity drivers across a given chemical series).
5. Overlay blocker scaffolds to our drain-plug model, and identify the BC, BP, and BY features. Increase the de-solvation cost (reflected in the polarity [33,34]) of these features to slow k_on_ (especially BY, when present), while maintaining on-target activity.
6. Attenuate the pKa of unavoidable basic nitrogen(s) in BP to slow k_on_ and collaterally minimize undesirable lysosomal trapping and membrane partitioning effects manifesting as high volume of distribution (V_ss_).
7. Maximally decrease percent inhibition at the projected TFPC_max_, which corresponds to the TTIC_max_ adjusted for lysosomal/membrane/off-target binding.
8. Assess pro-arrhythmicity based on Δ(exposure) between the projected TFPC_max_ and hERG IC_50_ (noting that *in vitro* potency may be overestimated for non-trappable blockers). The simulated general dose-response relationship for trappable and non-trappable blockers as a function of k_on_ and k_off_ (for non-trappables) and IC_50_ (for trappables) is shown in Figure 12.
9. hERG blockade in the absence of multiple ion channel effects (MICE) is pro-arrhythmic, as is blockade of other single cardiac ion channels. The emphasis by other workers on MICE [11] may reflect the poor translatability of *in vitro* hERG IC_50_ data to humans, although the normal inward-outward ion current balance is certainly disrupted by MICE (except in the case of verapamil, in which blockade of inward I_Ca_ is believed to offset blockade of outward I_Kr_).

**Figure 12.**
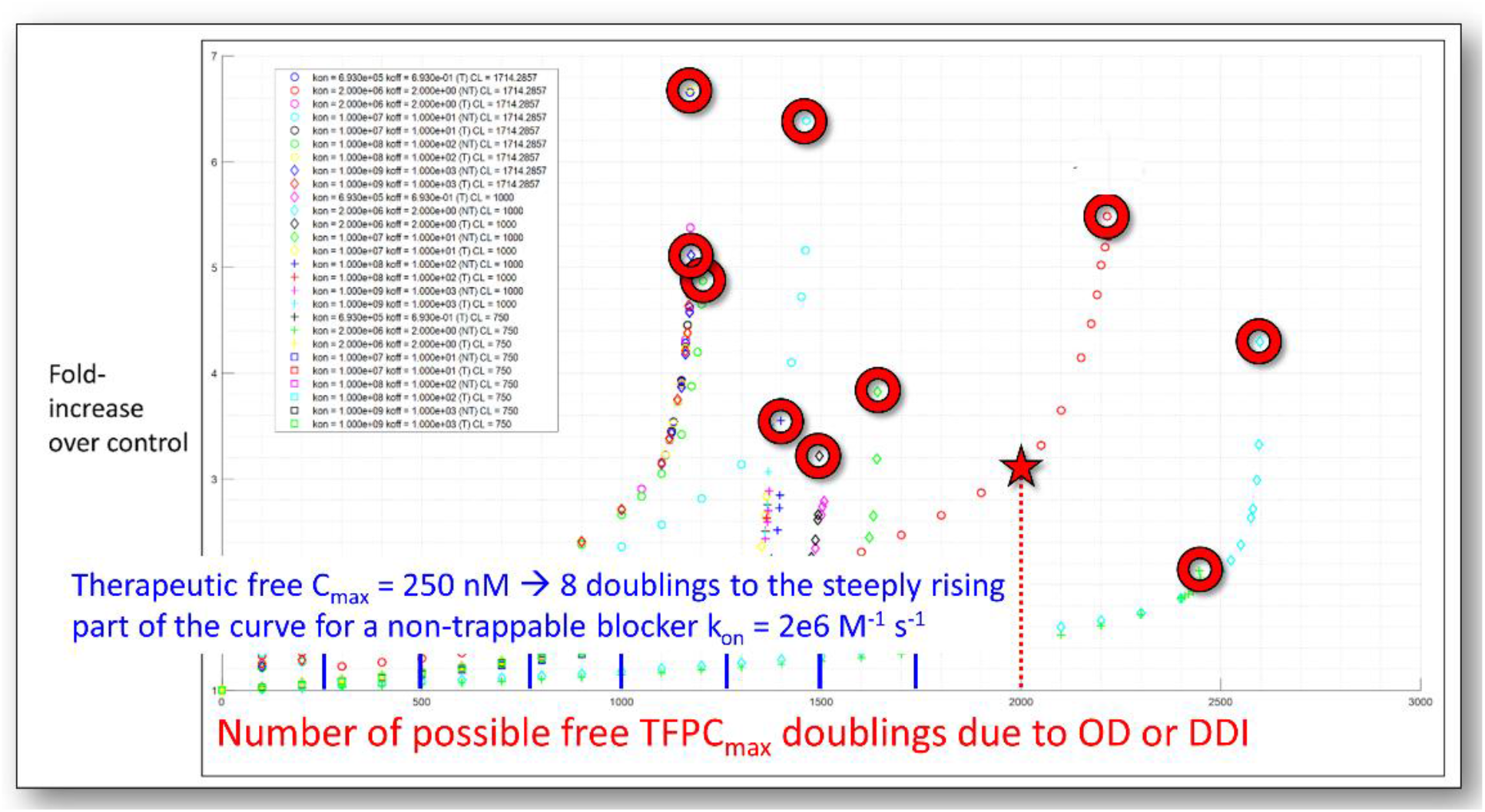
Hook-shaped dose-response curves for trappable and non-trappable blockers simulated using our modified version of the O’Hara-Rudy model of the undiseased human heart as a function of k_on_, k_off_, and hERG IC_50_. The hollow circles denote the “pre-arrhythmic” tipping-point exposure at the precipice of the arrhythmic response. The hERG safety margin should reside somewhere on the quasi-linear region of the corresponding curve (far from the tipping point) within a certain number of exposure doublings between the TFPC_max_ and tipping point, commensurate with the benefit/risk of the disease indication. The safety margin decreases with increasing percent inhibition at the TFPC_max_, but potentially more slowly than suggested by the Redfern SI.

### Chloroquine/hydroxychloroquine case study

Chloroquine and its hydroxy derivative (HCQ) are currently of interest as potential SARS-CoV-2 anti-viral therapies [42]. The anti-viral efficacy of these drugs is unconfirmed (and is under considerable scrutiny [43]), whereas both drugs are known to exhibit pro-arrhythmic and arrhythmic levels of hERG blockade in the clinical setting (based on data from the Federal Adverse Event Reporting System summarized in [44], a recent COVID-19 patient cohort [45], and reports of QT prolongation above 500 ms in COVID-19 patients [46]). Furthermore, hERG is likely blocked as well by the principal metabolites of HCQ, which consist of desethylchloroquine, desethylhydroxychloroquine, and bisdesethylchloroquine [47]. The reported hERG IC_50_s for chloroquine and HCQ are 2.5 μM [28] and ~5.6 μM (the latter of which we estimated from the Hill equation based on 35% inhibition at 3.0 μM reported in [29]). Both drugs fit unambiguously to our hERG blocker overlay model (shown for chloroquine in Figure 13), with BP terminating well below the putative constriction zone in the closed channel state in the absence of a BY group (consistent with trappability). We note that the putatively higher IC_50_ of HCQ is consistent with the expected greater incremental de-solvation cost of the hydroxyl group, which is predicted by our model to reside on BP. Neglecting the additive effects of azithromycin or other drug combinations, the maximum safe TFPC_max_ for chloroquine and HCQ under the Redfern SI equates to 83 and 187 nM (i.e. 2.5E-6/30 and 5.6E-6/30 μM), respectively. The mean total plasma concentration of <u>orally administered</u> HCQ in one COVID-19 study is 0.46 μg/ml [42], equating to ~1.4 μM (compared with ~1.2 μM in malaria patients [48]), which may or may not be representative of all patients. Confirmation of the reported potency and plasma PK of HCQ and its metabolites <u>at therapeutic COVID-19 dosing levels</u> is essential for accurate hERG safety assessment. The high V_ss_ of HCQ and its metabolites [48] results in extremely slow renal excretion (mean terminal t_1/2_ of 40-50 days), raising the possibility of multi-dose accumulation. The estimated TFPC_max_ of HCQ adjusted for an ~50% plasma protein bound fraction [47] is ~700 nM (i.e. 0.5 ? 1.4E-6 μM), or ~3.7-fold above the Redfern SI (i.e. 700/187). The extent to which the TI may be underestimated by the Redfern criterion depends on the true FIC_max_ in humans (noting that the fractional hERG occupancy of trappable blockers accumulates to FIC_max_/(FIC_max_ + hERG IC_50_). Chloroquine and HCQ lack definitive preclinical hERG <u>safety margins</u> vis-à-vis the Redfern criterion, which is exacerbated by putative hERG trappability and low plasma protein binding (possibly traded against high lysosomal uptake [49]). HCQ has nevertheless realized many prescription-years as an anti-malarial [50] and auto-immune therapy, the safety profile of which is well-understood for those specific indications [43]. All safety indices are context dependent, and can be exceeded (and the TI lost) in cases of significant exposure escalation above the established therapeutic level due to additive multi-drug effects, and/or the impact of underlying disease on drug clearance or the safety threshold. LQT monitoring and exposure control are essential for off-label HCQ administration, given:

1. That arrhythmia can result from even <u>transient</u> excursions in exposure ≥ the arrhythmic threshold. The high potential for cardiovascular impairment in COVID-19 patients may result in a downward shift of our predicted arrhythmic hERG occupancy level (which is based on the undiseased heart).
2. Co-administration with other pro-arrhythmic drugs, including azithromycin (hERG IC_50_ = 219 μM [51]) may lower the safe exposure level of HCQ (noting that reported cases of azithromycin-induced arrhythmia [44] have been attributed to intra-cellular Na^+^ loading [51]).
3. That HCQ metabolites likewise block hERG and exhibit long t_1/2_.
4. The high potential for impaired clearance due to renal compromise in COVID-19 patients.
5. The potential for DDIs in COVID-19 patients undergoing multi-drug therapies.
6. That intravenous HCQ administration results in up to 19-fold higher blood levels (2,436 ng/ml [48], translating to ~13 μM) compared with oral administration. We note that this level is more than 2-fold above our estimated hERG IC_50_.

**Figure 13.**
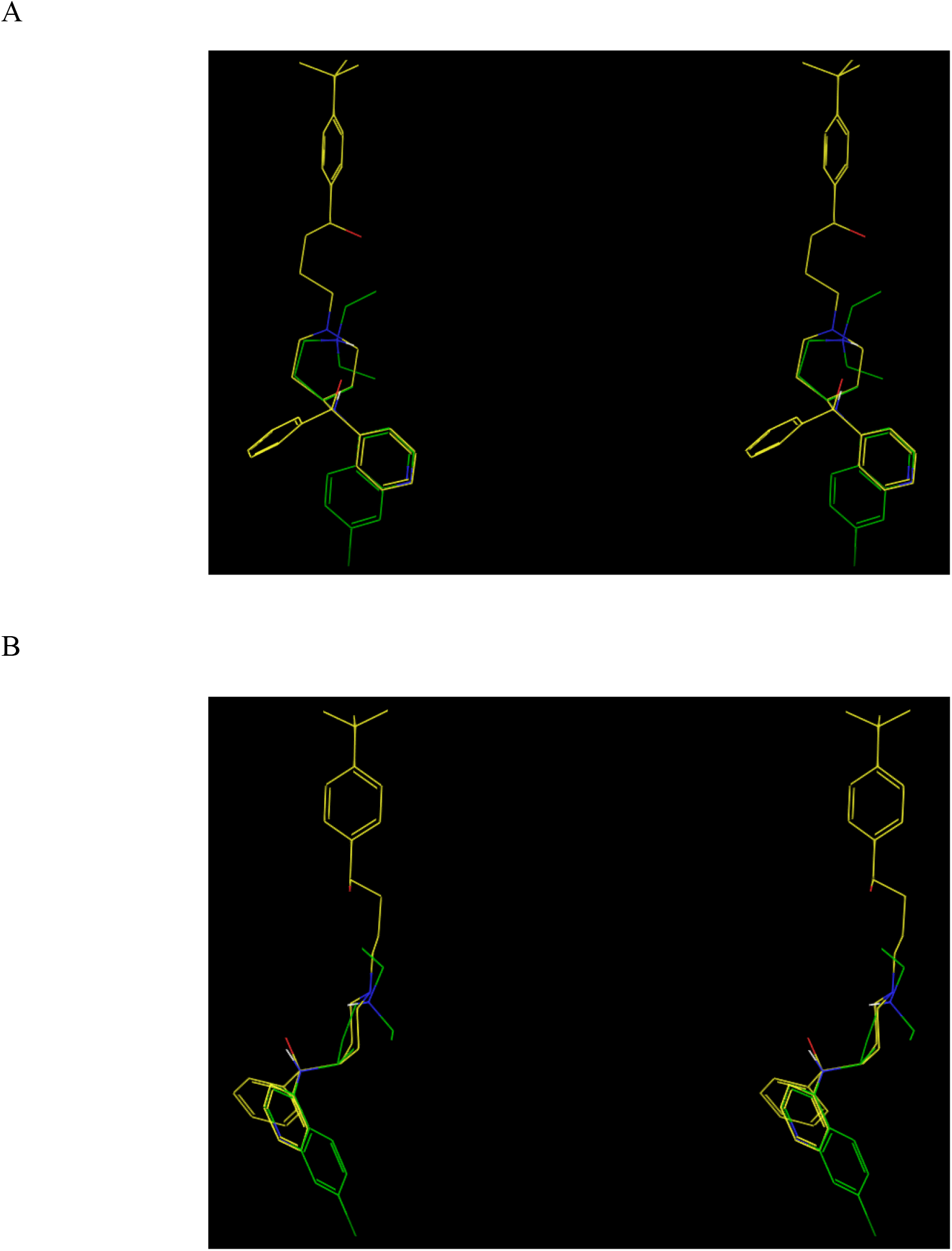
(A) Front view of chloroquine (green carbons) overlaid on our hERG blocker model (terfenadine shown for reference (yellow carbons)). (B) Side view of the overlay shown in A.

## Discussion

We used non-equilibrium structure-free energy (Biodynamics) principles, combined with a threedimensional ligand-based alignment of a set of trappable and non-trappable hERG blockers and published cryo-EM structures of hERG and Na_v_1.4 [26,27], to understand the non-equilibrium structure-free energy relationships governing blocker-hERG binding. Specifically, we propose that mutual blocker-hERG de-solvation and re-solvation costs that respectively govern k_on_ and k_off_ are localized to a set of blocker-specific docking interfaces (denoted as C-BC, P-BP, and Y-BY) in the C-linker and pore cavities. This hypothesis is consistent with our previous ligand-based analysis of hERG blocker chemical space (based on the Redfern dataset [39] and internal data), including neutral bisaryl, basic bisaryl, and alkylamine-containing scaffolds [17].

hERG safety assessment and mitigation are necessarily weighted toward the prevention of false negatives over false positives (erring on the side of caution), which is entirely justified given the acute, life-threatening implications of TdP, combined with:

1. Uncertainty in the true cause-effect relationship between measured hERG inhibition and arrhythmic risk.
2. The chicken-egg nature of hERG assessment/mitigation during the lead optimization stage, stemming from the lack of *in vivo* ECG and PK data needed to establish a human-relevant safety margin. Critical unknown parameters during this stage include:

a. The predicted TFPC_max_ in humans.
b. The predicted TFIC_max_ at the TFPC_max_ in humans, noting that percent hERG inhibition cannot be assessed from IC_50_s in the absence of TFIC_max_ information. Furthermore, TFIC_max_ may be influenced by potentially unquantifiable lysosomal trapping, membrane binding, primary target binding, and off-target binding.
c. Confirmed hERG IC_50_ and percent inhibition data, noting that such data is typically measured using less accurate high throughput techniques (Figure 11), and furthermore, may overestimate the <u>dynamic</u> occupancy of non-trappable blockers.

Simultaneous satisfaction of the highly stringent Redfern criterion (translating approximately to zero tolerated hERG inhibition at the TFPC_max_) and therapeutic target criteria is extremely challenging, time-consuming, and failure prone. The question is whether absolute hERG safety at all possible exposures, and across all indications and patient populations is an acceptable tradeoff against slowed progress and failure among hERG-afflicted R&D programs addressing unmet medical needs (which our findings can only help inform, but not answer).

We predicted arrhythmogenic propensity in terms of fractional occupancy of the ventricular hERG channel population in the <u>undiseased</u> human heart at exposures ranging between the TFIC_max_ and maximum FIC_max_ expected from DDIs or overdose (noting that the possible need for dose escalation during clinical trials must be considered in establishing the hERG SI). We define the human-relevant SI in terms of the Δ(exposure) between the true TFIC_max_ (typically > the efficacious free C_max_, allowing for metabolic clearance) and the arrhythmic exposure, which may be significantly less in the presence of cardiac dysfunction. The actual safety margin depends on the maximum Δ(exposure) due to PK variability, DDIs, overdose, or dose escalation (Figure 14). In contrast, the Redfern SI begins with near zero tolerated hERG inhibition at the TFPC_max_, translating to a 100% safety margin relative to the putative arrhythmic inhibition level. Furthermore, the Redfern SI appears biased toward potent trappable blockers, which based on our previous analysis [17], demand a greater safety margin than non-trappables.

**Figure 14.**
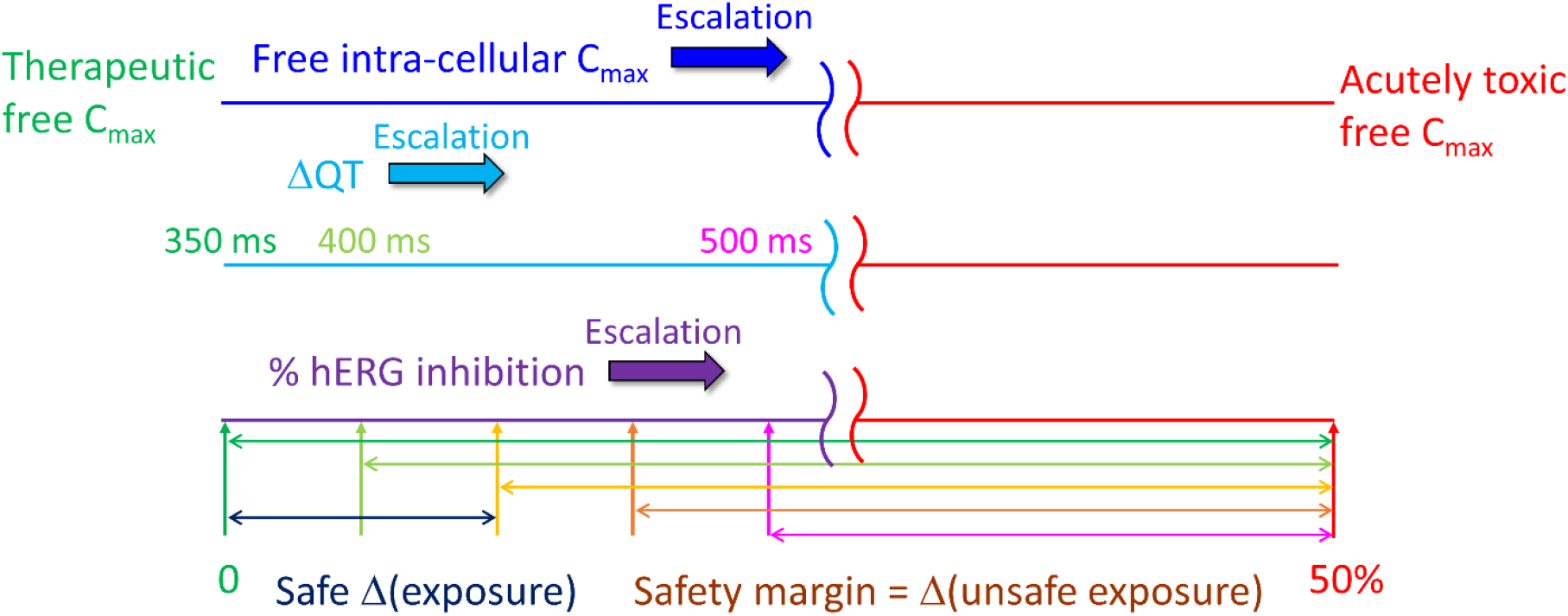
Our proposed safety margin is defined in terms of the fractional hERG occupancy at the <u>therapeutic</u> intra-cellular free C_max_ relative to that at the predicted <u>arrhythmic</u> exposure level in the otherwise normal heart. This metric differs from the Redfern SI, in which hERG percent inhibition is restricted to effectively zero at the TFPC_max_ (corresponding to the bright green double-headed arrow in the lower part of the figure). The safe Δ(exposure) depends on the benefit/risk of the disease indication, but should always correspond to hERG occupancy far below the arrhythmic level (assumed here as 50% of the open/conducting channel population [17]), and for all practical purposes, QT < 500 ms.

## Conclusions

In this work, we emphasize the need for human-relevant hERG safety prediction and mitigation criteria during the preclinical stages of drug discovery, accounting for the true relationships between chemical structure and *in vivo*-relevant dynamic hERG occupancy, and between dynamic occupancy and the pro-arrhythmic effects thereof on the <u>otherwise normal</u> human cardiac AP. Both relationships seem to be poorly understood under the conventional wisdom (including reliance on static equilibrium binding metrics and near zero tolerance for blockade at therapeutic exposures). We propose that blocker association is driven by steric shape complementarity, blocker desolvation cost at key docking interfaces (together with electrostatic attraction in the case of basic compounds), and dissociation is driven largely by mutual blocker and binding site re-solvation costs (noting that the dissociation rate of non-trappable blockers under physiological conditions is necessarily faster than the rate of channel closing). We further propose a drain-plug-like canonical binding mode, in which blockers straddle the pore and C-linker cavities (analogous to the binding mode of GDN in the cryo-EM structure of Na_v_1.4 [27]), projecting R-groups into non-bulk-like solvation sites in the C-linker (C/BC), and pore (P/BP, Y/BY) regions of the outer vestibule. We attribute trappability to a putative constriction zone that forms during channel closing, with which only planar blocker groups (or the absence of groups in this position) are sterically compatible. Disruption of blocker binding is putatively achievable via trappable ➔ non-trappable chemical transformations, together with incorporation of polar groups at BC, BP, and BY, thereby increasing the blocker de-solvation cost at those sites. We showed that the Redfern SI equates effectively to zero hERG occupancy at the TFPC_max_, and is weighted heavily toward the prevention of false negative blockers from entering clinical trials at the possible expense of false positives. Our approach, guided by accurately measured *in vitro* percent inhibition and human-relevant *in vivo* PK data, may help to minimize trial-and-error mitigation, and lower uncertainty in human-relevant hERG safety predictions.

## Acknowledgements

We thank Dr. Laszlo Urban and Dr. Anatoli Lvov for providing the hERG radio-ligand binding and Qpatch data shown in Figure 11.

## References

1. Sanguinetti MC, Tristani-Firouzi M. hERG potassium channels and cardiac arrhythmia. Nature. 2006;440: 463–469.

2. Curran ME, Splawski I, Timothy KW, Vincen GM, Green ED, Keating MT. A molecular basis for cardiac arrhythmia: HERG mutations cause long QT syndrome. Cell. 1995;80: 795–803. doi:10.1016/0092-8674(95)90358-5

3. Roden DM. Long QT syndrome: Reduced repolarization reserve and the genetic link. J Intern Med. 2006;259: 59–69. doi:10.1111/j.1365-2796.2005.01589.x

4. Sanguinetti MC, Jiang C, Curran ME, Keating MT. A mechanistic link between an inherited and an acquird cardiac arrthytmia: HERG encodes the IKr potassium channel. Cell. 1995;81: 299–307. doi:10.1016/0092-8674(95)90340-2

5. Haverkamp W, Breithardt G, Camm AJ, Janse MJ, Rosen MR, Antzelevitch C, et al. The Potential for QT Prolongation and Proarrhythmia by Non-antiarrhythmic Drugs: Clinical and Regulatory Implications, Sophia Antipolis, France, 24-25 June 1999. Eur Heart J. 2000;21: 1216–1231. doi:10.1053/euhj.2000.2249

6. Viskin S. Long QT syndromes and torsade de pointes. Lancet. 1999;354: 1625–1633. doi:10.1016/S0140-6736(99)02107-8

7. Liang SI, van Lengerich B, Eichel K, Cha M, Patterson DM, Yoon TY, et al. Phosphorylated EGFR Dimers Are Not Sufficient to Activate Ras. Cell Rep. 2018;22: 2593–2600. doi:10.1016/j.celrep.2018.02.031

8. Keating MT, Sanguinetti MC. Molecular and cellular mechanisms of cardiac arrhythmias. Cell. 2001;104: 569–580. doi:10.1016/B978-0-12-381510-1.00041-7

9. Kalyaanamoorthy S, Barakat KH. Development of Safe Drugs: The hERG Challenge. Med Res Rev. 2018;38: 525–555. doi:10.1002/med.21445

10. Siramshetty VB, Nickel J, Omieczynski C, Gohlke BO, Drwal MN, Preissner R. WITHDRAWN - A resource for withdrawn and discontinued drugs. Nucleic Acids Res. 2016;44: D1080–D1086. doi:10.1093/nar/gkv1192

11. Colatsky T, Fermini B, Gintant G, Pierson JB, Sager P, Sekino Y, et al. The Comprehensive in Vitro Proarrhythmia Assay (CiPA) initiative — Update on progress. J Pharmacol Toxicol Methods. The Authors; 2016;81: 15–20. doi:10.1016/j.vascn.2016.06.002

12. Redfern WS, Carlsson L, Davis AS, Lynch WG, MacKenzie I, Palethorpe S, et al. Relationships between preclinical cardiac electrophysiology, clinical QT interval prolongation and torsade de pointes for a broad range of drugs: Evidence for a provisional safety margin in drug development. Cardiovasc Res. 2003;58: 32–45. doi:10.1016/S0008-6363(02)00846-5

13. Kirsch GE, Trepakova ES, Brimecombe JC, Sidach SS, Erickson HD, Kochan MC, et al. Variability in the measurement of hERG potassium channel inhibition: Effects of temperature and stimulus pattern. J Pharmacol Toxicol Methods. 2004;50: 93–101. doi:10.1016/j.vascn.2004.06.003

14. Winiowska B, Polak S. HERG in vitro interchange factorsdevelopment and verification hERG in vitro interchange factors Barbara Winiowska and Sebastian Polak. Toxicol Mech Methods. 2009;19: 278–284. doi:10.1080/15376510902777194

15. Windley MJ, Lee W, Vandenberg JI, Hill AP. The temperature dependence of kinetics associated with drug block of hERG channels is compound-specific and an important factor for proarrhythmic risk prediction. Mol Pharmacol. 2018;94: 760–769. doi:10.1124/mol.117.111534

16. Rudy Y, Hara TO. Simulation of the Undiseased Human Cardiac Ventricular Action Potential: Model Formulation and Experimental Validation. 2011;7. doi:10.1371/journal.pcbi.1002061

17. Pearlstein RA, Andrew MacCannell K, Erdemli G, Yeola S, Helmlinger G, Hu Q-Y, et al. Implications of Dynamic Occupancy, Binding Kinetics, and Channel Gating Kinetics for hERG Blocker Safety Assessment and Mitigation. Curr Top Med Chem. 2016;16: 1792–1818. doi:10.2174/1568026616666160315142156

18. Selvaggio G, Pearlstein RA. Biodynamics: A novel quasi-first principles theory on the fundamental mechanisms of cellular function/dysfunction and the pharmacological modulation thereof. PLoS One. 2018;13: e0202376. doi:10.1371/journal.pone.0202376

19. Pearlstein RA, Hu Q-Y, Zhou J, Yowe D, Levell J, Dale B, et al. New hypotheses about the structure-function of proprotein convertase subtilisin/kexin type 9: Analys is of the epidermal growth factor-like repeat A docking site using watermap. Proteins Struct Funct Bioinforma. 2010;78: 2571–2586. doi:10.1002/prot.22767

20. Pearlstein RA, Sherman W, Abel R. Contributions of water transfer energy to proteinligand association and dissociation barriers: Watermap analysis of a series of p38α MAP kinase inhibitors. Proteins Struct Funct Bioinforma. 2013;81. doi:10.1002/prot.24276

21. Tran Q-T, Pearlstein RA, Williams S, Reilly J, Krucker T, Erdemli G. Structure-kinetic relationship of carbapenem antibacterials permeating through E. coli OmpC porin. Proteins Struct Funct Bioinforma. 2014;82. doi:10.1002/prot.24659

22. Tran QT, Williams S, Farid R, Erdemli G, Pearlstein R. The translocation kinetics of antibiotics through porin OmpC: Insights from structure-based solvation mapping using WaterMap. Proteins Struct Funct Bioinforma. 2013;81: 291–299. doi:10.1002/prot.24185

23. Pearlstein RA, McKay DJJ, Hornak V, Dickson C, Golosov A, Harrison T, et al. Building New Bridges between In Vitro and In Vivo in Early Drug Discovery: Where Molecular Modeling Meets Systems Biology. Curr Top Med Chem. 2017;17: 1–1. doi:10.2174/1568026617666170414152311

24. Farid R, Day T, Friesner RA, Pearlstein RA. New insights about HERG blockade obtained from protein modeling, potential energy mapping, and docking studies. Bioorganic Med Chem. 2006;14: 3160–3173. doi:10.1016/j.bmc.2005.12.032

25. Windisch A, Timin EN, Schwarz T, Stork-Riedler D, Erker T, Ecker GF, et al. Trapping and dissociation of propafenone derivatives in HERG channels. Br J Pharmacol. 2011;162: 1542–1552. doi:10.1111/j.1476-5381.2010.01159.x

26. Wang W, MacKinnon R. Cryo-EM Structure of the Open Human Ether-à-go-go-Related K+ Channel hERG. Cell. Elsevier Inc.; 2017;169: 422–430.e10. doi:10.1016/j.cell.2017.03.048

27. Pan X, Li Z, Zhou Q, Shen H, Wu K, Huang X, et al. Structure of the human voltagegated sodium channel Nav1.4 in complex with β1. Science (80-). 2018;362: eaau2486. doi:10.1126/science.aau2486

28. Traebert M, Dumotier B, Meister L, Hoffmann P, Dominguez-Estevez M, Suter W. Inhibition of hERG K+ currents by antimalarial drugs in stably transfected HEK293 cells. Eur J Pharmacol. Elsevier; 2004;484: 41–48. doi:10.1016/j.ejphar.2003.11.003

29. Capel RA, Herring N, Kalla M, Yavari A, Mirams GR, Douglas G, et al. Hydroxychloroquine reduces heart rate by modulating the hyperpolarization-activated current If: Novel electrophysiological insights and therapeutic potential. Hear Rhythm. Elsevier B.V.; 2015;12: 2186–2194. doi:10.1016/j.hrthm.2015.05.027

30. Dickson CJ, Velez-Vega C, Duca JS. Revealing Molecular Determinants of hERG Blocker and Activator Binding. Cite This J Chem Inf Model. 2019;2020: 192–203. doi:10.1021/acs.jcim.9b00773

31. Recanatini M, Cavalli A, Masetti M. Modeling hERG and its interactions with drugs: Recent advances in light of current potassium channel simulations. ChemMedChem. 2008;3: 523–535. doi:10.1002/cmdc.200700264

32. Cavalli A, Poluzzi E, De Ponti F, Recanatini M. Toward a Pharmacophore for Drugs Inducing the Long QT Syndrome: Insights from a CoMFA Study of HERG K + Channel Blockers. 2002; doi:10.1021/jm0208875

33. Nagatomo T, Kawakami K, Kikuchi K, Takemasa H, Oginosawa Y, Tsurugi T, et al. Poster P3-6: A Comparison of hERG channel blocking activities by beta-blockers - implication for clinical strategy.

34. Cheung AK, Hurley B, Kerrigan R, Shu L, Chin DN, Shen Y, et al. Discovery of Small Molecule Splicing Modulators of Survival Motor Neuron-2 (SMN2) for the Treatment of Spinal Muscular Atrophy (SMA). 2018; doi:10.1021/acs.jmedchem.8b01291

35. Barber MJ, Wendt DJ, Starmer CF, Grant AO. Blockade of cardiac sodium channels: Competition between the permeant ion and antiarrhythmic drugs. J Clin Invest. 1992;90: 368–381. doi:10.1172/JCI115871

36. Stork D, Timin EN, Berjukow S, Huber C, Hohaus a, Auer M, et al. State dependent dissociation of HERG channel inhibitors. Br J Pharmacol. 2007;151: 1368–1376. doi:10.1038/sj.bjp.0707356

37. Yeh JZ, Armstrong CM. Immobilisation of gating charge by a substance that simulates inactivation. Nature. 1978;273: 387–389. doi:10.1038/273387a0

38. Mitcheson JS, Chen J, Sanguinetti MC. Trapping of a methanesulfonanilide by closure of the HERG potassium channel activation gate. J Gen Physiol. 2000;115: 229–240. doi:10.1085/jgp.115.3.229

39. Redfern WS, Carlsson L, Davis AS, Lynch WG, MacKenzie I, Palethorpe S, et al. Relationships between preclinical cardiac electrophysiology, clinical QT interval prolongation and torsade de pointes for a broad range of drugs: Evidence for a provisional safety margin in drug development. Cardiovascular Research. 2003. pp. 32–45. doi:10.1016/S0008-6363(02)00846-5

40. Yu Z, IJzerman AP, Heitman LH. Kv11.1 (hERG)-induced cardiotoxicity: A molecular insight from a binding kinetics study of prototypical Kv11.1 (hERG) inhibitors. Br J Pharmacol. 2015;172: 940–955. doi:10.1111/bph.12967

41. Morotti S, Polak S, Li ZhihuaLi Z, Chang KC, Dutta S, Mirams GR, et al. Uncertainty Quantification Reveals the Importance of Data Variability and Experimental Design Considerations for in Silico Proarrhythmia Risk Assessment. Front Physiol | www.frontiersin.org. 2017;8: 917. doi:10.3389/fphys.2017.00917

42. Gautret P, Lagier J-C, Parola P, Hoang VT, Meddeb L, Mailhe M, et al. Hydroxychloroquine and azithromycin as a treatment of COVID-19: results of an openlabel non-randomized clinical trial. Int J Antimicrob Agents. Elsevier BV; 2020; 105949. doi:10.1016/j.ijantimicag.2020.105949

43. Mehra MR, Desai SS, Ruschitzka F, Patel AN. Articles Hydroxychloroquine or chloroquine with or without a macrolide for treatment of COVID-19: a multinational registry analysis. Lancet. 2020; doi:10.1016/S0140-6736(20)31180-6

44. Giudicessi JR, Noseworthy PA, Friedman PA, Ackerman MJ. Mayo Foundation for Medical Education and Research. Mayo Clin Proc. Mayo Clinic Proceedings; 2020.

45. Mercuro NJ, Yen CF, Shim DJ, Maher TR, McCoy CM, Zimetbaum PJ, et al. Risk of QT Interval Prolongation Associated With Use of Hydroxychloroquine With or Without Concomitant Azithromycin Among Hospitalized Patients Testing Positive for Coronavirus Disease 2019 (COVID-19). JAMA Cardiol. 2020; doi:10.1001/jamacardio.2020.1834

46. Giudicessi JR, Roden DM, Wilde AAM, Ackerman MJ. Journal Pre-proof Genetic Susceptibility for COVID-19-Associated Sudden Cardiac Death in African Americans. 2020; doi:10.1016/j.hrthm.2020.04.045

47. Lim HS, Im JS, Cho JY, Bae KS, Klein TA, Yeom JS, et al. Pharmacokinetics of hydroxychloroquine and its clinical implications in chemoprophylaxis against malaria caused by plasmodium vivax. Antimicrob Agents Chemother. 2009;53: 1468–1475. doi:10.1128/AAC.00339-08

48. fda, cder. PLAQUENIL ® HYDROXYCHLOROQUINE SULFATE TABLETS, USP DESCRIPTION [Internet]. Available: http://www.cdc.gov/malaria

49. Al-Bari AA. Chloroquine analogues in drug discovery: New directions of uses, mechanisms of actions and toxic manifestations from malaria to multifarious diseases. J Antimicrob Chemother. 2014;70: 1608–1621. doi:10.1093/jac/dkv018

50. Lim HS, Im JS, Cho JY, Bae KS, Klein TA, Yeom JS, et al. Pharmacokinetics of hydroxychloroquine and its clinical implications in chemoprophylaxis against malaria caused by plasmodium vivax. Antimicrob Agents Chemother. 2009;53: 1468–1475. doi:10.1128/AAC.00339-08

51. Yang Z, Prinsen JK, Bersell KR, Shen W, Yermalitskaya L, Sidorova T, et al. Azithromycin Causes a Novel Proarrhythmic Syndrome. Circ Arrhythmia Electrophysiol. 2017;10. doi:10.1161/CIRCEP.115.003560

